# Three neurotransmitters regulate diverse inhibitory and excitatory Parvalbumin interneuron circuits in the dorsal horn

**DOI:** 10.1101/2020.12.23.424233

**Authors:** MA Gradwell, KA Boyle, TJ Browne, AC Dickie, AM Bell, J Leonardo, FS Peralta Reyes, KM Smith, RJ Callister, CV Dayas, DI Hughes, BA Graham

## Abstract

Parvalbumin-expressing interneurons (PVINs) in the spinal dorsal horn are found primarily in laminae II inner and III. Inhibitory PVINs (iPVINs) play an important in segregating innocuous tactile input from pain-processing circuits, achieved through presynaptic inhibition of myelinated low-threshold mechanoreceptors and postsynaptic inhibition of distinct spinal circuits. By comparison, relatively little is known of the role of excitatory PVINs (ePVINs) in sensory processing. Here we use neuroanatomical and optogenetic approaches to show that ePVINs comprise a larger proportion of the PVIN population than previously reported, and that both ePVIN and iPVIN populations form synaptic connections amongst (and between) themselves. We find that these cells contribute to neuronal networks that influence activity within several functionally distinct circuits, and that aberrant activity of ePVINs under pathological conditions contributes to the development of mechanical hypersensitivity.

## Introduction

The dorsal horn of the spinal cord plays a key role in gating and modulating sensory input originating from primary afferents before it is relayed to supraspinal sites for perception. The principal neuronal populations in this region can be differentiated into interneurons that process and modulate sensory input at a local (spinal) level, and projection neurons that relay this information to higher centres for sensory perception. Interneurons can be further subdivided into two groups based on their principal neurotransmitter content: excitatory interneurons use glutamate, whereas inhibitory interneurons use GABA and/or glycine (Todd, 2010). Both populations of interneurons are highly heterogeneous in terms of their morphology, physiological properties, neurochemistry, and genetic profiles. Recent advances in molecular-genetic profiling of dorsal horn interneurons (Abraira et al., 2017; Sathyamurthy et al., 2018; Häring et al., 2018) coupled the increased use of transgenic animals have enabled researchers to target and manipulate discrete neuronal populations with great precision (Graham and Hughes, 2019), leading to a better understanding of the functional role of various cell types (Peirs and Seal, 2016; Abraira et al., 2017; Moehring et al., 2018).

One population of spinal interneurons that have been widely studied are those that express the calcium-binding protein parvalbumin (PV). In the dorsal horn, these cells are found primarily in lamina II inner (IIi) and III, with similar patterns of expression being described in various species including rat (Yamamoto et al., 1989; Celio, 1990; Antal et al., 1990), cat (Anelli and Heckman, 2005) and mouse (Hughes et al., 2012). Immunohistochemical studies in the rat have shown that ∼75% of PV-immunolabelled cells in laminae I-III express both GABA and glycine (Antal et al., 1991; Laing et al., 1994), with the remaining cells considered to be excitatory (glutamatergic) interneurons. More recent approaches using transgenic mouse lines have estimated that 70% of genetically-labelled PVINs are inhibitory interneurons, with the remainder being excitatory (Abraira et al., 2017). Inhibitory PVINs (iPVINs) are known to be important in setting mechanical thresholds under both normal and pathological conditions (Hughes et al., 2012; Petitjean et al., 2015; Boyle et al., 2019). Under normal conditions, these cells mediate both presynaptic (axoaxonic) inhibition of myelinated low-threshold mechanoreceptive afferents (A-LTMRs) and postsynaptic inhibition of both vertical cells (Boyle et al., 2019) and PKCγ-expressing interneurons (Petitjean et al., 2015). Following nerve injury, the intrinsic excitability of iPVINs is reduced and the resulting disinhibition opens circuits through which A-LTMR input can be relayed to lamina I. These studies demonstrate the importance of inhibition mediated by iPVINs under normal and pathological conditions, but the role of excitatory PVINs in spinal circuits has yet to be determined. To address this, the aim of our study was to characterise the synaptic connectivity of both excitatory and inhibitory PVINs, and define the dorsal horn circuits that allow these populations influence the relay of sensory information to supraspinal sites.

## Results

### Molecular-genetic profiling of PV neurons in laminae IIi and III

We have previously shown that the distribution of PV-expressing neurons in the mouse spinal dorsal horn is similar to that found in other species with most cells found in laminae IIi and III (Hughes et al., 2012), but note that the incidence of these cells in the mouse dorsal horn is higher than in either rat or cat (Yamamoto et al., 1989; Celio, 1990; Antal et al., 1990; Anelli and Heckman, 2005). Despite this difference, immunohistochemical approaches in the rat (Laing et al., 1994) and molecular-genetic studies in the mouse (Abraira et al., 2017) estimate that 70-75% of PVINs are inhibitory interneurons, with the remainder being excitatory interneurons. We aimed to determine the relative proportions of excitatory and inhibitory interneurons within the PV population more directly by using a multiple-labelling fluorescent *in situ* hybridization approach (Figure 1). In these studies, parvalbumin-expressing cells were identified using a probe for PValb, excitatory interneurons using a probe for Slc17a6 (to identify VGLUT2-expressing cells), and inhibitory interneurons using a probe for GAD1 (to identify GABAergic cells). The distribution of cells labelled with the PValb probe matched that reported previously using immunohistochemistry (Hughes et al., 2012), with most cells being found in lamina IIi and III (Figure 1A). From a total of 515 PV-expressing cells analysed in sections from three mice (147, 153 and 215), we found that 52.7% (SEM ± 4.7 %) expressed Slc17a6 (274/515: 64, 90 and 120), with the remainder expressing GAD1 (241/515; 83, 63 and 95; Figure 1B).

**Figure 1.**
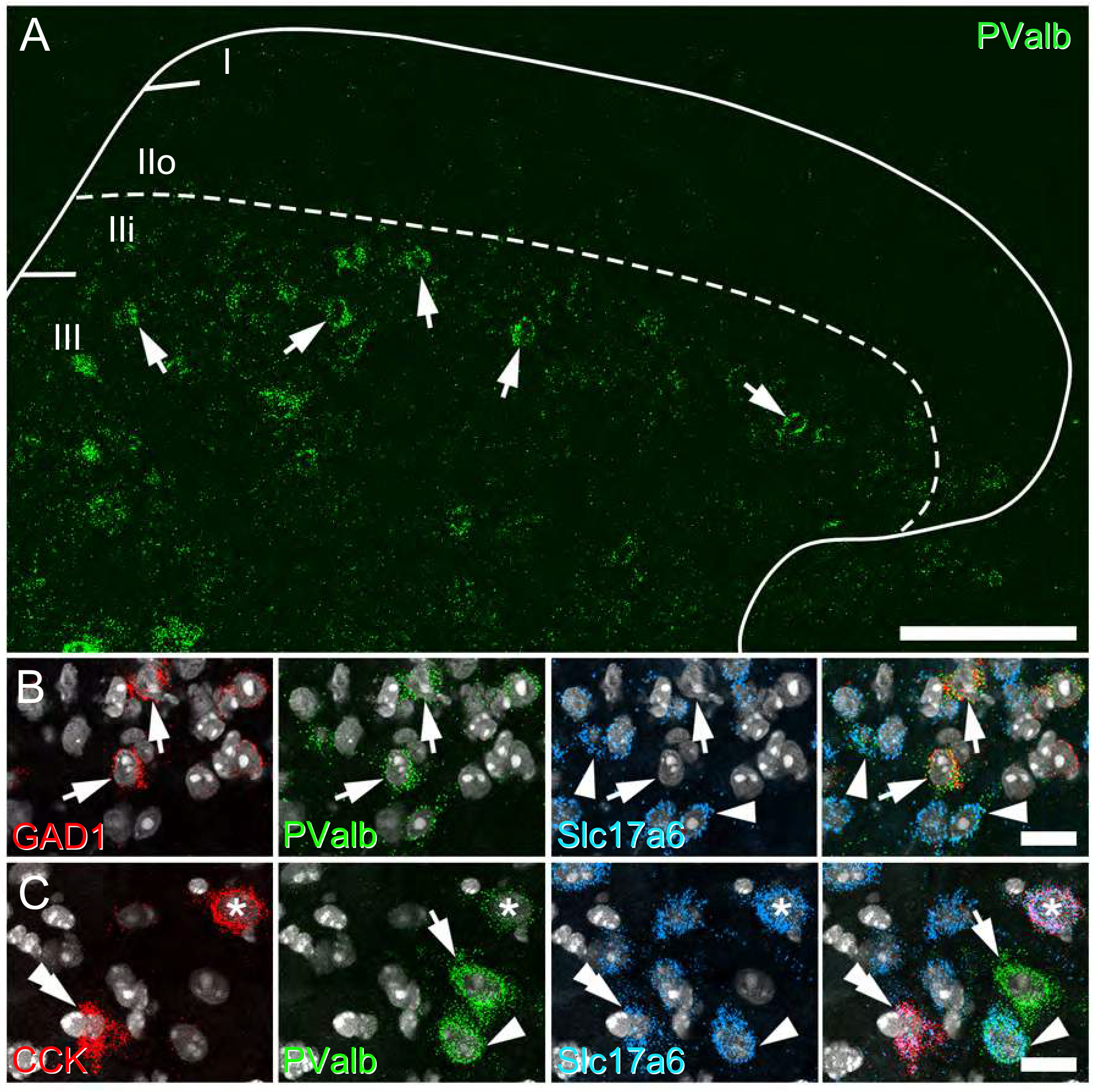
Neurochemical characterisation of parvalbumin cells in the spinal dorsal horn using fluorescent in situ hybridisation. **A**, Lumbar spinal cord sections processed for fluorescent *in situ* hybridisation to map parvalbumin expression (PValb; green) showed most cells were concentrated in laminae IIi and III (arrows). **B**, Multiple labelling with probes for GAD1 (red; inhibitory interneurons), PValb (green), Slc17ac (blue; for excitatory interneurons) and the NucBlue (grey) was used to show that approximately half of the PVINs in laminae I-III were inhibitory interneurons (arrows), with the remainder being excitatory interneurons (arrowheads). **C**, Similar studies using probes to CCK (red), PValb (green), Slc17a6 (blue) and NucBlue (grey) to show that approximately 75% of excitatory PVINs co-express CCK (asterisk), but these account for only ∼25% of CCK population (double arrowhead). This field also shows an ePVIN (arrowhead) and and iPVIN (arrow), neither of which co-express CCK. All images are generated from a single optical section. Scale bars (µm): A = 100; B and C = 25.

Molecular-genetic profiling studies of mouse dorsal horn interneurons have also noted PV expression in both glutamatergic (Glut1 and Glut2) and inhibitory (Gaba14 and Gaba15) subtypes (Häring et al., 2018), and we have used this dataset to further define the neurochemical properties of ePVINs. Both Glut1 and Glut2 subtypes described in this work were shown to express high levels of the neuropeptide CCK, and these cells are largely restricted to lamina III, where PVINs are most common (Cortes et al., 1990; Abelson and Micevych, 1991; Schiffmann et al, 1991; Abraira et al., 2017). We therefore aimed to determine the degree of co-expression of CCK in the ePVIN populations (Figure 1C). Out of 201 Slc17a6-expressing PV cells analysed from three mice (75, 54 and 72), 74.5% (± 2.3%) expressed CCK (150; 59, 40 and 51). Conversely, from a total of 556 Slc17a6 cells that express CCK in these laminae, only 27.0% express PV (59, 40 and 51). From these studies, we conclude that ePVINs account for a far higher proportion of PVINs than previously reported, and while most of these (∼75%) express CCK, though ePVINs only account for approximately a quarter of CCK-expressing cells in this region.

We have previously used offspring from a PV^Cre^ mouse line crossed with a Cre-dependent reporter line (Ai9) to target and record from iPVINs in the spinal dorsal horn (Boyle et al., 2019). The fidelity of tdTomato (tdTOM) expression in PV-expressing cells from these PV^Cre^;Ai9 mice was established by comparing the incidence of reporter protein in cells labelled with antibodies raised against PV in the spinal cord (dorsal horn laminae II and III, and in the ventral horn), the cerebellum, and both the dentate gyrus and CA1 subfield the hippocampus (Supplementary Figure S1). Overall, the vast majority of tdTOM-expressing cells co-expressed immunolabelling for PV (94.5%, 1504 out of 1591 cells), and the majority of PV-immunoreactive (PV-IR) cells also expressed tdTOM (68.2%; 1504 out of 2204 cells). A high proportion of PV-IR cells in the hippocampus (83.1%; ±0.7) and ventral horn of the spinal cord (86.3%; ±7.9) expressed tdTOM, and nearly all tdTOM-labelled cells in these regions were immunolabelled for PV (94.0%; ±0.9 and 97.7%; ± 0.7, hippocampus and ventral horn, respectively). By contrast, far fewer PV-IR cells in dorsal horn laminae II and III expressed tdTOM (37%; ± 10.1), although the vast majority of these tdTOM cells were immunolabelled for PV (90.4%; ±2.2). We therefore find that Cre-mediated recombination in this PV^Cre^ line captures PV-expressing cells with high fidelity, but that the overall proportion of PV-IR cells labelled with tdTOM is lower in the dorsal horn than other regions of the central nervous system.

Given that we have used this mouse line to record from and label iPVINs, as well as to manipulate their function using optogenetic and targeted silencing approaches (Boyle et al., 2019), we determined whether tdTOM expression in these dorsal laminae captured iPVINs preferentially by comparing the expression pattern of a developmental marker for inhibitory interneurons, Pax2 as previously (Smith et al., 2015). We found that 48.0 % (± 2.1) of PV-IR cells in lamina II and III showed immunolabelling for Pax2, whereas 40.2% (± 1.4) of cells that co-express both tdTOM and PV-IR were also immunolabelled for Pax2. These immunohistochemical findings mirror our *in situ* hybridization analysis above and show that there is no preferential expression of PV^Cre^-mediated reporter molecules in in iPVIN and ePVIN populations of lamina II and III. We conclude that this line provides a reliable, unbiased means of studying PVINs. The resultant expression profile provides a faithful means to resolve detailed neuroanatomical and functional connectivity patterns within the complex and heterogenous neuropil of the spinal dorsal horn.

### Morphological features and interconnectivity of PVINs in laminae IIi and III

We have previously reconstructed the morphology of individual PVINs from both tissue sections immunolabelled for PV and of Neurobiotin-filled cells from targeted whole-cell patch-clamp recording in slices from PV^Cre^;Ai9 mice (Hughes et al., 2012; Boyle et al., 2019). Cells reconstructed from immunolabelled tissue displayed various morphologies (Hughes et al., 2012), whereas those reconstructed following patch-clamp recording and identified as inhibitory interneurons were typically islet cells (Boyle et al., 2019). While targeted recordings of neurons from slices and Neurobiotin-recovery morphological reconstruction of recorded cells with great confidence, it is technically demanding and throughput of cells for morphological analyses is low. This approach does, however, lend itself to unintended bias in cell selection for recordings, potentially skewing sampling towards larger cells (Smith et al., 2015). Furthermore, recording conditions for slices maintained *in vitro* and fixation protocols for subsequent anatomical studies can have important bearing on the integrity of the recovered morphology as well as the preservation of tissue ultrastructure and antigenicity. We therefore aimed to compare the morphology of ePVINs and iPVINs in a more systematic manner using optimal experimental conditions.

This analysis adopted an approach that utilises Brainbow-labelling of PVINs through intraspinal injections of Cre-dependent viral vectors and optimal fixation protocols to preserve tissue structure and antigenicity (Cai et al., 2013). Individual PVINs expressed unique colour profiles based on the stochastic expression of farnesylated fluorescent proteins encoded by the respective viruses, while both the cellular and subcellular localisation of various target molecules remained unaltered (Figure 2A). We assessed the morphological features of individual Brainbow-labelled ePVINs and iPVINs from confocal image stacks, as differentiated by expression (or absence) of the inhibitory cell marker Pax2 (Figure 2B). The somatodendritic arborisation of 64 Brainbow-labelled neurons in laminae IIi-III were traced in 3 dimensions using Neurolucida for confocal software (MBF Bioscience, VT, USA). Thirty of these cells expressed Pax2 and were classified as iPVINs, with the remainder classified as ePVINs. Although some overlap in the morphological features was evident between the two populations, iPVINs generally had larger cell bodies, dendritic arbors that extended further in the rostrocaudal axis and branched more often, compared to ePVINs (Figure 2C, D, E). The soma volumes of iPVINs were significantly greater than those of ePVINs (841.8 μm^3^ *vs*. 444.3μm^3^, respectively, p < 0.0001), as were their total dendritic lengths (2054 μm *vs*. 943 μm, p < 0.0001). The difference in dendritic length between these two populations was most apparent when measured in the rostro-caudal (RC) axis (1603µm *vs*. 696µm for iPVINs and ePVINs, respectively, p < 0.0001), with iPVINs also showing greater dendritic spread in the RC axis (433.4 µm *vs*. 236.2 µm, p < 0.0001). The volume of tissue encapsulated by the dendritic arbors of iPVINs was also significantly larger than that of ePVINs (660654 μm^3^ to 144623 μm^3^, respectively, p < 0.0001), while fractal dimension (1.12 *vs*. 1.08, p = 0.0139) and tortuosity of dendrites within these volumes (42.3 *vs*. 20.7, p < 0.0001) were also significantly higher for iPVINs than ePVINs.

**Figure 2.**
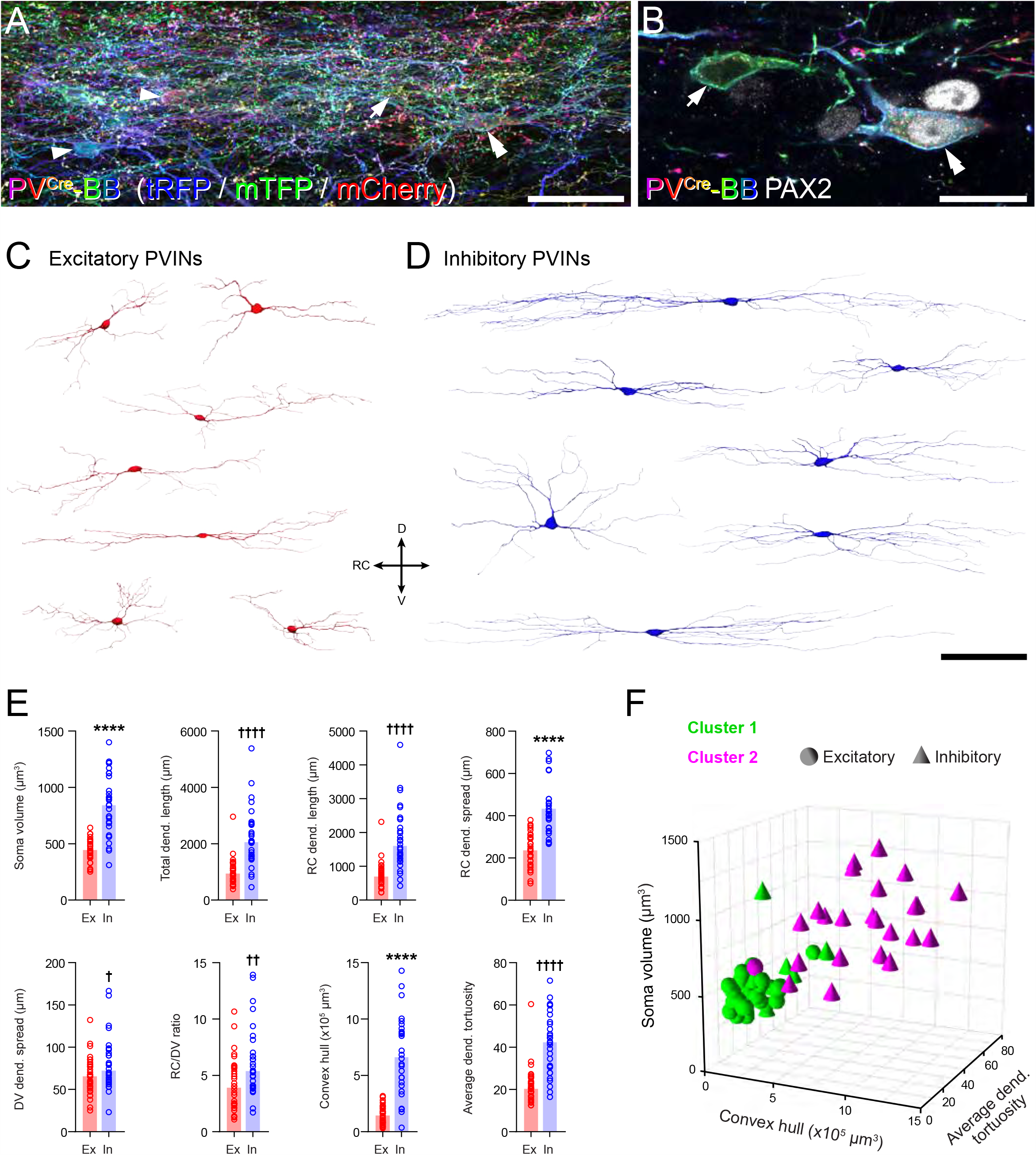
Morphometric analyses of Brainbow-labelled excitatory and inhibitory PV interneurons. **A**, Example of Brainbow labelling in laminae IIi/III of a sagittal section from a PV^Cre^ mouse injected with AAV.Brainbow1 and AAV.Brainbow2, showing a dense plexus of Brainbow-labelled PV cells in laminae IIi and III. Examples of individual cell bodies within this plexus are highlighted with arrowheads. The cells highlighted with an arrow and a double arrowhead are shown at higher magnification in **B**. This image is a maximum projection of 107 optical sections at 0.5 µm z-spacing. **B**, Higher magnification of two cells outlined in panel A (arrow and double arrowhead), generated from a single optical section. This figure demonstrates the use of immunostaining for Pax2 to identify inhibitory (double arrowhead; Pax2-expressing) and excitatory (arrow; lacking Pax2) Brainbow-labelled PVINs. **C and D**, Representative 3D reconstructions of the somato-dendritic morphology of excitatory (C, red) and inhibitory (D, blue) Brainbow-labelled PVINs. DV = dorsoventral axis, RC = rostro-caudal axis. **E**, Grouped scatterplots of selected morphometric parameters of all reconstructed excitatory (Ex; red; n = 34) and inhibitory (In; blue; n = 30) PVINs. Key to y-axes: Dend = dendritic; RC dend. length = total length of dendrite projecting in the rostro-caudal axis; RC dend. spread and DV dend. spread are the distances between the most distal points in the rostro-caudal and dorsoventral axes, respectively; RC/DV ratio = RC dend. spread/DV dend. spread. **** = p<0.0001 by unpaired t-test for normally distributed data; † = p<0.05, †† = p<0.01, †††† = p<0.0001 by Mann-Whitney test for non-normally distributed data. Bars for normally distributed data show mean, bars for non-normally distributed data show median. **F**, Scatterplot of soma volume (y-axis) versus convex hull volume (x-axis) versus average dendritic tortuosity for all reconstructed PV interneurons (z-axis), grouped by k means-derived cluster (Cluster 1 or Cluster 2; green or magenta, respectively) and neurotransmitter phenotype (excitatory or inhibitory; spheres or cones, respectively). Note that these three axes variables were selected for this visualisation as they are assumed to be independent of each other. Scale bars (µm): A = 50; B = 20; C and D = 100.

In order to determine whether morphological features can be used to differentiate both PVIN subpopulations, we performed K-means multivariate cluster analysis on fifty morphological parameters extracted from the Neurolucida reconstructions of these cells. These comprised 5 parameters for the soma and 45 for the dendritic arbors (including those plotted in Figure 2E; see Table 1 for details). By setting the number of clusters to 2, and the distribution of both PVIN subpopulations within these K-means-derived clusters could be compared in an unbiased manner (Figure 2F and Table 1). Using this approach, 97 % (33/34) of ePVINs were contained within Cluster 1 and 77% (23/30) of iPVINs were within Cluster 2. From this, we conclude that excitatory and inhibitory PVINs are morphologically distinct, and that soma size and extent of dendritic arbors in the rostro-caudal axis are two of the strongest discriminators between these populations.

**Table 1.** Distribution of excitatory and inhibitory PV interneurons within K-means-derived clusters. Contingency table showing the count of excitatory and inhibitory PVINs within each of the two K-means-derived clusters (Cluster 1 and Cluster 2).

|  | Cluster 1 | Cluster 2 | Total |
| --- | --- | --- | --- |
| Excitatory | 33 | 1 | **34** |
| Inhibitory | 7 | 23 | **30** |
| **Total** | **40** | **24** | **64** |

Having prepared this tissue to optimise structural preservation and the retention of tissue antigenicity, we aimed to determine whether these PVIN subpopulations formed synaptic connections to each other. Immunolabelling for Pax2 was again used to differentiate iPVINs from ePVINs, and once this had been established, we were able to trace axons from these cells to determine whether they formed homotypic and/or heterotypic synaptic connections on to other Brainbow-labelled PVINs. Sparce labelling afforded in the PV^Cre^ line made it possible to trace individual axons with greater precision than would otherwise be possible. Excitatory and inhibitory synapses were visualised using immunolabelling for Homer-1 and gephyrin, respectively. Using these approaches, we found anatomical evidence for excitatory synaptic inputs derived from ePVINs onto both iPVINs and ePVINs (Figures 3A and 3B, respectively), and also of inhibitory synaptic inputs from iPVINs onto both iPVINS and ePVINS (Figures 3C and 3D, respectively). We conclude that the synaptic targets of both excitatory and inhibitory PVINs include other PVINs in laminae IIi and III, and that these cells form both homotypic and heterotypic connections.

**Figure 3.**
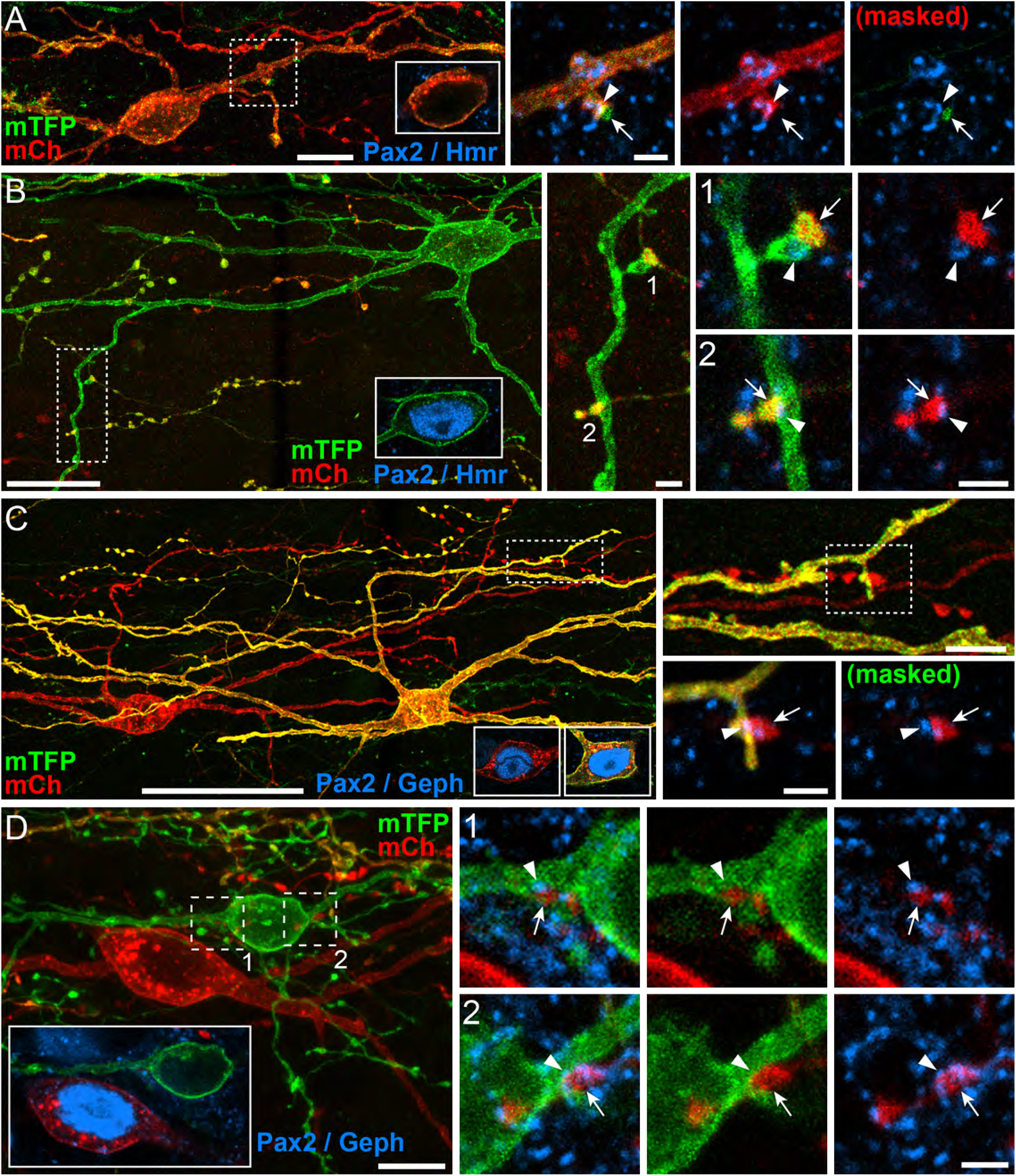
Homotypic and heterotypic synaptic connectivity between PVINs in laminae IIi and III. **A and B**, Examples of homotypic synaptic connections made by ePVINs on to other ePVINs (**A**), and heterotypic synaptic connections on to iPVINs (**B**). Insets in A and B show presence or absence Pax2 (blue) in the soma of the target neuron. High power insets of areas outlined on the target neurons show axon terminals (arrows) forming excitatory synaptic inputs on to the dendrites of excitatory (**A**) and inhibitory PVINs (**B**), respectively. Excitatory synapses are verified by the presence of immunolabelling for Homer1 (blue; arrowheads). **C and D**, Examples of homotypic synaptic connections made by iPVINs on to other iPVINs (**C**), and heterotypic synaptic connections on to ePVINs (**D**). Insets in C and D show presence or absence Pax2 (blue) in soma of the pre-and postsynaptic neurons illustrated. High power insets of areas outlined on the target neurons show axon terminals (arrows) forming inhibitory synaptic inputs on to the dendrites of inhibitory (**C**) and excitatory PVINs (**D**), respectively. Inhibitory synapses are verified by the presence of immunolabelling for gephyrin (blue; arrowheads). Lower power panels are maximum projections of 35, 115, 132 and 47 optical sections for figures A, B, C and D, respectively, with z-separation of 0.5 µm (**A** and **C**) or 0.3 µm (**B** and **D**). Insets detailing Pax2-immunolabelling in cell bodies are single optical sections. High power panels detailing synaptic contacts are maximum projections generated from 3 optical sections at 0.3µm z-steps. Scale bars (µm): A = 10 and 2; B = 20, 2 and 2; C = 50, 5 and 2; D = 10 and 2.

### PVIN photostimulation

To study the function of PVIN-mediated synaptic connections we bred PV^Cre^;Ai32 mice to undertake channelrhodopsin2 (ChR2)-assisted circuit mapping. We first assessed the expression of YFP (corresponding to ChR2 expression) immunolabelling in the dorsal horn of spinal cord sections of these animals. The distribution of YFP immunolabelling mirrored that of PV-IR, with a dense plexus of labelling in lamina IIi and III, but largely absent in more dorsal lamina (Figure 4A). The majority of YFP/ChR2^+^ neurons were also immunolabelled for PV (98%, 133/136 and 106/109 in 2 animals). However, consistent with sparse dorsal horn expression in the PV^Cre^ mouse line, only a subset of PV-IR neurons expressed ChR2 (∼10%, 133/1420 and 106/1110 in 2 animals). Therefore, while this mouse line only captures a proportion of the entire PV population, it reliably and selectively expresses ChR2 in these cells. As highlighted above for neuroanatomical work, relatively sparse ChR2 expression among PVINs permitted detailed optogenetic analysis of connectivity within the dorsal horn. This restricted expression patterns allows the photostimlation responses to be studied with greater precision from individual neurons, and avoids the problem of widespread network activity that would be evident in lines that capture a higher yield of cells.

**Figure 4.**
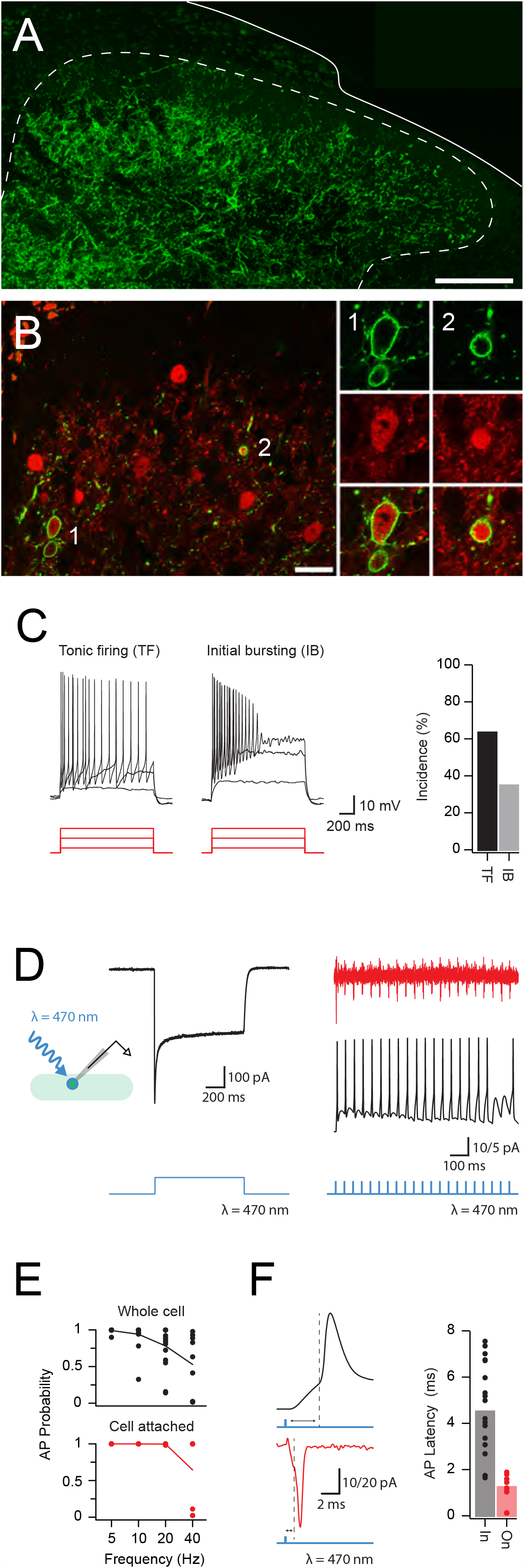
ChR2 expression and activation in PVINs. **A**, Representative image showing the distribution of ChR2:YFP expression (green) in the lumbar dorsal horn of a PV^Cre^;Ai32 mouse. **B**, Left panel compares ChR2:YFP expression (green) and PV-IR profiles (red). The majority of ChR2:YFP neurons express PV-IR (98%) examples noted (1, 2), though relatively few PV-IR profiles express ChR2:YFP (9%), double arrow. Right panels show neurons labelled ‘1’ and ‘2’ in right panel at high magnification; ChR2:YFP (upper), PV-IR (middle), merge (lower). **C**, Upper traces show action potential discharge in PVINs recorded during depolarising step current injections (lower, 20, 60, 100 pA steps shown). PVIN discharge patterns could be reliably classified as either tonic firing, or initial bursting. Bar plot (right) shows incidence of PVIN discharge patterns. **D**, Trace shows example photocurrent recording from a PVIN. Recorded in voltage-clamp, PVINs exhibit large inward photocurrents in response to photostimulation. Inset schematic shows recording arrangement with photostimulation. **E**, Traces show 1ms photostimulation at 20Hz reliably evokes AP discharge in PVINs in cell-attached voltage clamp recordings (upper, red) and whole cell current clamp recording (lower, black). Blue trace (bottom) indicates photostimulation protocol for each representative trace. Plots (right) compare reliability of evoked AP discharge across a range of photostimulation frequencies using whole-cell (upper) and cell-attached (lower) recording configurations. **F**, Traces compare photostimulation-evoked action potential spiking using whole-cell (upper) and cell-attached (lower) recording configurations. Note, recruitment latency (time between photostimulation and AP threshold) is shorter in the cell-attached mode. Bar graphs (right) show variability in recruitment latency using whole-cell (black) and cell-attached (red) recording configurations. Short latencies observed in the cell-attached configuration suggest that the ChR2-expressing PVINs require at least ∼2 ms to generate an AP during photostimulation. Scale bar in A = 200 μm.

Targeted recordings assessing action potential discharge in ChR2/YFP-expressing PVINs during depolarising current step injections (Figure 4B) showed that these neurons exhibited either tonic firing (n = 34/53, persistent AP discharge throughout the depolarizing step) or initial bursting responses (n = 19/53, AP discharge limited to depolarizing step onset). These discharge patterns match our previous work in PVeGFP mice and PV^Cre^;Ai9 mice (Hughes et al., 2012; Abraira et al., 2017; Boyle et al., 2019), and suggest that the ChR2/YFP-expressing cells captured by the PV^Cre^;Ai32 line are representative of the PVIN population. Photostimulation of recorded PVINs produced immediate large inward currents (photocurrents) in voltage clamp (holding potential −70 mV) and these photocurrents were sufficient to evoke AP discharge in current clamp mode (Figure 4B-C). The capacity of PVINs to fire repetitively during photostimulation was also examined by varying the frequency of brief photostimulation pulses (1 ms duration). APs could be reliably evoked by photostimulation at 5 Hz (99% whole cell *vs*. 100% cell attached), however, the probability of successful AP generation decreased at higher frequencies (10 Hz: 94% whole cell *vs*. 100% on cell; 20 Hz: 78% whole cell *vs*. 100% cell attached; 40Hz: 53% whole cell *vs*. 65% cell attached). The mean latency from photostimulation onset to AP discharge, or recruitment delay, in PVINs was 4.6 ± 0.5 ms in the whole-cell recording mode, and 1.3 ± 0.5 ms during cell-attached recordings (Figure 4D). These data confirm ChR2 expression is sufficient in cells from the PV^Cre^;Ai32 cross to optically generate spiking in PVINs at frequencies of 20 Hz (in cell attached recordings) with a delay of 1.3 ms after photostimulation.

### PVINs provide monosynaptic input onto neighbouring PVINs and other dorsal horn populations

The postsynaptic targets of PVINs in laminae I, II outer (IIo), II inner (IIi) and III were assessed by recording from unidentified dorsal horn neurons (cells lacking YFP in laminae I-III), and PVINs (YFP-expressing cells in laminae II-III). Recordings from unidentified neurons (UN) were further subdivide into those located within the plexus of PVIN dendrites and processes (UN laminae IIi-III), and those located dorsal to this region (UN laminae I-IIo). Given that PVINs have predominantly been studied in the context of inhibitory connections, we first used a CsCl based internal to accentuate inhibitory currents. Photostimulation often elicited optically-evoked postsynaptic currents (oPSCs, Figure 5A), which occurred at short latencies (3.9 ± 0.1 ms *vs*. 4.1 ± 0.3 ms *vs*. 4.9± 0.2 ms for UN:LIIi-III, UN:LI-IIo, and PVINs, respectively) and with limited jitter (0.6 ± 0.1 ms *vs*. 0.7 ± 0.1 ms *vs*. 0.4 ± 0.1 ms; UN:LIIi-III, UN:LI-IIo, and PVINs, respectively). Allowing for the recruitment delay of PVIN photostimulation (1.3 ms, cell attached) and conduction plus synaptic delays of ∼2ms (Lu and Perl, 2003), we conclude that these latencies and limited jitter are consistent with the existence of monosynaptic connections. Using these criteria, PVIN photostimulation resulted in monosynaptic oPSCs in 79% (146/185) of UN:LIIi-III, 30% (16/53) of UN:LI-IIo, and 61% (67/110) of PVIN recordings in laminae II-III.

**Figure 5.**
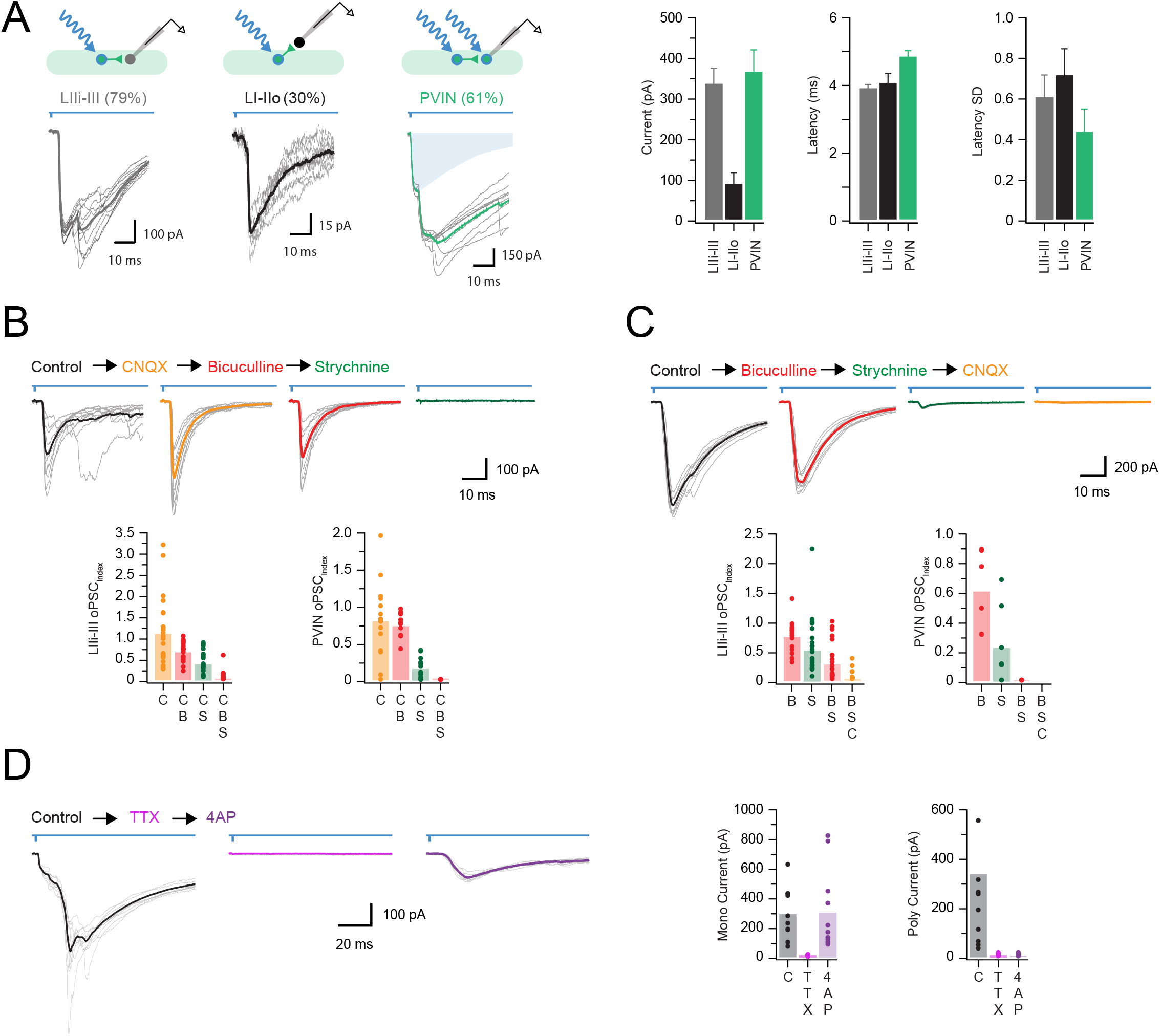
iPVINs provide mixed, glycine dominant postsynaptic inhibition. **A**, Upper schematics show the three recording configurations used to study monosynaptic connections: PVIN to lamina IIi-III neurons, PVIN to lamina I-IIo neurons and PVIN to PVIN. Voltage clamp recordings (−70 mV) showing optically-evoked postsynaptic currents (oPSCs) in laminae I-IIo and IIi-III neurons, as well as in PVINs themselves (10 consecutive sweeps and average overlayed). Blue shading in PVIN trace denotes underlying photocurrent isolated by pharmacological block of synaptic events. Group data to right compares oPSC amplitude, latency, and latency standard deviation (jitter), suggesting oPSCs have monosynaptic characteristics. **B**, Representative oPSCs recorded from a single neuron (black), and following sequential bath application of CNQX (orange), bicuculline (red), and strychnine (green). Group data below shows effects of this drug application approach in lamina IIi-III neurons (left), and PVINs (right), highlighting glycine dominant oPSCs (i.e. the PSC index is most reduced by strychnine). **C**, Representative oPSCs recorded from a single neuron (black) and following sequential addition of bicuculline (red), strychnine (green), and CNQX (orange). Note the small oPSC remaining following addition of bicuculline and strychnine, suggesting a glutamatergic component. Group data below shows effects of drug application in lamina II_i_-III neurons (left), and PVINs (right). **D**, Representative oPSCs recorded from a single neuron (black) and following sequential bath application of TTX (pink) and 4AP (purple). Addition of TTX blocks action potential-dependent oPSCs, which can be recovered by enhancing excitability of axon terminals with 4AP. Group data below shows effects of drug application on monosynaptic currents (bottom left) and longer latency polysynaptic currents (bottom right), highlighting the drug cocktail’s ability to isolate monosynaptic oPSCs.

Group data comparisons (Figure 5A) show that oPSC amplitude was similar for both UN:LIIi-III and PVINs (both located within the PV plexus), but oPSC amplitude was significantly smaller in recordings from UN:LI-IIo (337 ± 39 pA *vs*. 367 ± 54 pA *vs*. 90 ± 28 pA, respectively, p < 0.05). These data suggest that within laminae IIi and III, PVINs form an extensive network of synaptic connections both with other PVINs as well as many other cells in this region. This is of particular interest given that recent genetic ablation and tetanus toxin silencing studies (Petitjean et al. 2015; Boyle et al. 2019) indicate these circuits play a critical role in the confinement of tactile signals to deeper layers of the dorsal horn, interpreted through specific inhibitory connections with vertical cells and cells identified by PKCγ-expression.

### Varied pharmacology of PVIN mediated monosynaptic inputs

The pharmacology of monosynaptic oPSCs was next assessed, as both excitatory and inhibitory currents could contribute to responses recorded using a CsCl-based internal. An oPSC_index_, calculated as the ratio of oPSC amplitude after antagonist application relative to oPSC amplitude under control conditions, was used to assess the contribution of different neurotransmitters to the oPSC response, with a values of 1 indicating no effect and 0 indicating complete block. Using this approach, bath addition of the AMPA/kainate receptor antagonist CNQX had variable effects on oPSCs (Figure 5B) ranging from complete block (oPSC_index_ = 0) to a marked increase (oPSC_index_ = 3.22). Overall, our group data suggests AMPA receptor block had limited affect oPSC amplitude in either UN:LIIi-III cells or PVINs (oPSC_index_: 1.13 ± 1.7, p = 0.43 and 0.82 ± 0.14, p = 0.21, respectively) but did reduce monosynaptic oPSC amplitude in recordings from UN:LI-IIo cells by half (oPSC_index_: 0.49 ± 0.15, p = 0.015). Thus, oPCSs in PVINs and UN:LIIi-III cells were largely insensitive to CNQX reflecting a high incidence of inhibitory oPSCs (oIPSCs), whereas a higher prevalence of CNQX-sensitive oPSCs in UN:LI-IIo cells (4/7) provides evidence for ePVIN-mediated excitatory oPSCs (oEPSC) in these superficial laminae.

Following isolation of monosynaptic inhibitory currents (oIPSCs) in CNQX, the pharmacology of these connections was further assessed by bath addition of the GABA_A_R antagonist bicuculline or the GlyR antagonist strychnine (Figure 5B). For recordings form UN:LIIi-III, UN:LI-IIo, and PVINs, bicuculline application decreased oIPSC amplitude (oIPSC_index_: 0.70 ± 0.05, p < 0.001, 0.49 ± 0.17, p = 0.2, and 0.75 ± 0.05, p = 0.001 for UN:LIIi-III, UN:LI-IIo, and PVINs). Similarly, bath application of strychnine also reduced oIPSC amplitude, though more dramatically than bicuculline (IPSC_index_: 0.42 ± 0.06, p < 0.001, 0.51 ± 0.19, p = 0.12, and 0.17 ± 0.04, p < 0.001 for UN:LIIi-III, UN:LI-IIo, and PVINs). Due to the varied effects of AMPA receptor block (Figure 5B) drug order was also reversed to better differentiate any role for glutamatergic signalling in short latency PVIN-mediated oPSCs (Figure 5C). Initial addition of bicuculline (PSC_index_: 0.77 ± 0.05, p < 0.001, 0.84 ± 0.07, p = 0.051, and 0.61 ± 0.11, p = 0.018 for UN:LIIi-III, UN:LI-IIo, and PVINs) and strychnine (PSC_index_: 0.53 ± 0.08, p < 0.001, 0.88 ± 0.14, p = 0.416, and 0.23 ± 0.10, p < 0.001 for UN:LIIi-III, UN:LI-IIo, and PVINs) reduced oPSC amplitude similar to above. Furthermore, co-application of bicuculline and strychnine abolished all responses in PVINs, but failed to completely block oPSCs in UN:LIIi-III cells (18/25) and UN:LI-IIo cells (8/8), respectively (oPSC_index_: 0.31 ± 0.06, p < 0.001 and 0.70 ± 0.16, p = 0.100). In these recordings, the remaining current was blocked by the bath addition of CNQX (oIPSC_index_: 0.06 ± 0.02, p < 0.001 and 0.14 ± 0.04, p < 0.001). These data confirm the existence of ePVIN-mediated oEPSCs in unidentified neuron located throughout LI-III.

The assignment of photostimulation responses as monosynaptic was also confirmed with bath application of TTX and 4-AP (Figure 5D), where only AP-independent ‘terminal release’ is possible by direct ChR2-mediated depolarization of presynaptic terminals (Petreanu et al. 2009, Cruikshank et al. 2010). TTX alone abolished all monosynaptic responses (i.e. those with short latency and low jitter), as well as any presumptive polysynaptic responses. In agreement with the criteria for monosynaptic responses, addition of 4-AP (in the presence of TTX) reinstated short latency responses (298 ± 56 pA *vs*. 10 ± 1 pA *vs*. 306 ± 82 pA; baseline *vs*. TTX *vs*. TTX + 4-AP, respectively). Finally, 4-AP reinstated monosynaptic oPSC amplitude was reduced by bicuculline (oIPSC_index_: 0.73 ± 0.07, p = 0.003 and 0.68 ± 0.8, p = 0.002 for UN:LIIi-III and PVINs), however, strychnine reduced oPSC amplitude more dramatically (oPSC_index_: 0.34 ± 0.06, p < 0.001 and 0.28 ± 0.7, p < 0.001 for UN:LIIi-III and PVINs). In 5/10 recordings from UN:LIIi-III cells, bicuculline and strychnine did not completely abolish 4-AP recovered oPSCs (oPSC_index_: 0.14 ± 0.04, p < 0.001). In these cases, the addition of CNQX abolished the remaining response (oIPSC_index_: 0.05 ± 0.02, p < 0.001). Taken together, these data confirm that both GABA and glycine are released by iPVINs with glycine being the dominant neurotransmitter in postsynaptic inhibition. Importantly, these data also identify the presence of ePVIN-mediated glutamatergic inputs within the DH. This matches the existence of a substantial excitatory PVIN population in our molecular profiling and Brainbow experiments.

### PVIN-mediated monosynaptic glutamatergic input rarely recruits other DH populations

To further examine the impact of ePVIN input within the dorsal horn, we next used a Kgluc-based internal solution to better differentiate excitatory and inhibitory responses and to study action potential spiking responses in postsynaptic targets. First, photostimulation of PVINs evoked monosynaptic oEPSCs in UN:LIIi-III (45% - 103/231), UN:LI-IIo (35% - 22/63), but not in PVINs (0% - 0/36, Figure 6A). Importantly, oEPSCs exhibited short latency (4.0 ± 0.1 ms and 3.3 ± 0.2 ms, for UN:LIIi-III and UN:LI-IIo) and limited jitter (0.7 ± 0.1 ms and 0.6 ± 0.1 ms for UN:LIIi-III and UN:LI-IIo), consistent with monosynaptic connections. oEPSC amplitude was similar in UN:LIIi-III and UN:LI-IIo cells (47.9 ± 5.6 pA, and 58.4 ± 23.0 pA, respectively). The capacity of these excitatory inputs to recruit action potential discharge was assessed in current clamp in some recordings (Figure 6B). This showed that oEPSPs were rarely capable of eliciting AP discharge in postsynaptic UN:LIIi-III (2/66) or UN:LI-IIo cells (2/23). To confirm the excitatory nature of these connections we assessed their sensitivity to bicuculine, strychnine, and CNQX application (Figure 6C). oEPSC amplitude for UN:LIIi-III and UN:LI-IIo was not reduced by bicuculline (oPSC_index_: 1.0 ± 0.04, p = 0.91 and 0.99 ± 0.09, p = 0.948, respectively) or strychnine (oPSC_index_: 0.90 ± 0.09, p = 0.31 and 1.83 ± 0.57, p = 0.201), but was abolished by CNQX (oEPSC_index_: 0.12 ± 0.01, p < 0.001 and 0.14 ± 0.03, p < 0.001). Finally, TTX was applied in a subset of recordings and abolished oEPSCs in all cases. These oEPSC could be restored, or in some cases enhanced, by the addition of 200 μM 4AP in TTX (oEPSC_index_: 3.7 ± 1.72, p = 0.138; Figure 6D), providing further evidence for the monosynaptic nature of these inputs. Consistent with the above pharmacology, the 4AP-ehanced currents were not affected by addition of bicuculline (oEPSC_index_: 1.03 ± 0.08, p = 0.732) or strychnine (oEPSC_index_: 0.97 ± 0.08, p = 0.786), but were abolished by CNQX (oEPSC_index_: 0.14 ± 0.05, p = 0.035). Taken together, these data show a subpopulation of glutamatergic PVINs provide excitatory drive to neurons in laminae IIi-III and I-IIo, however, these inputs rarely cause AP discharge in their postsynaptic targets.

**Figure 6.**
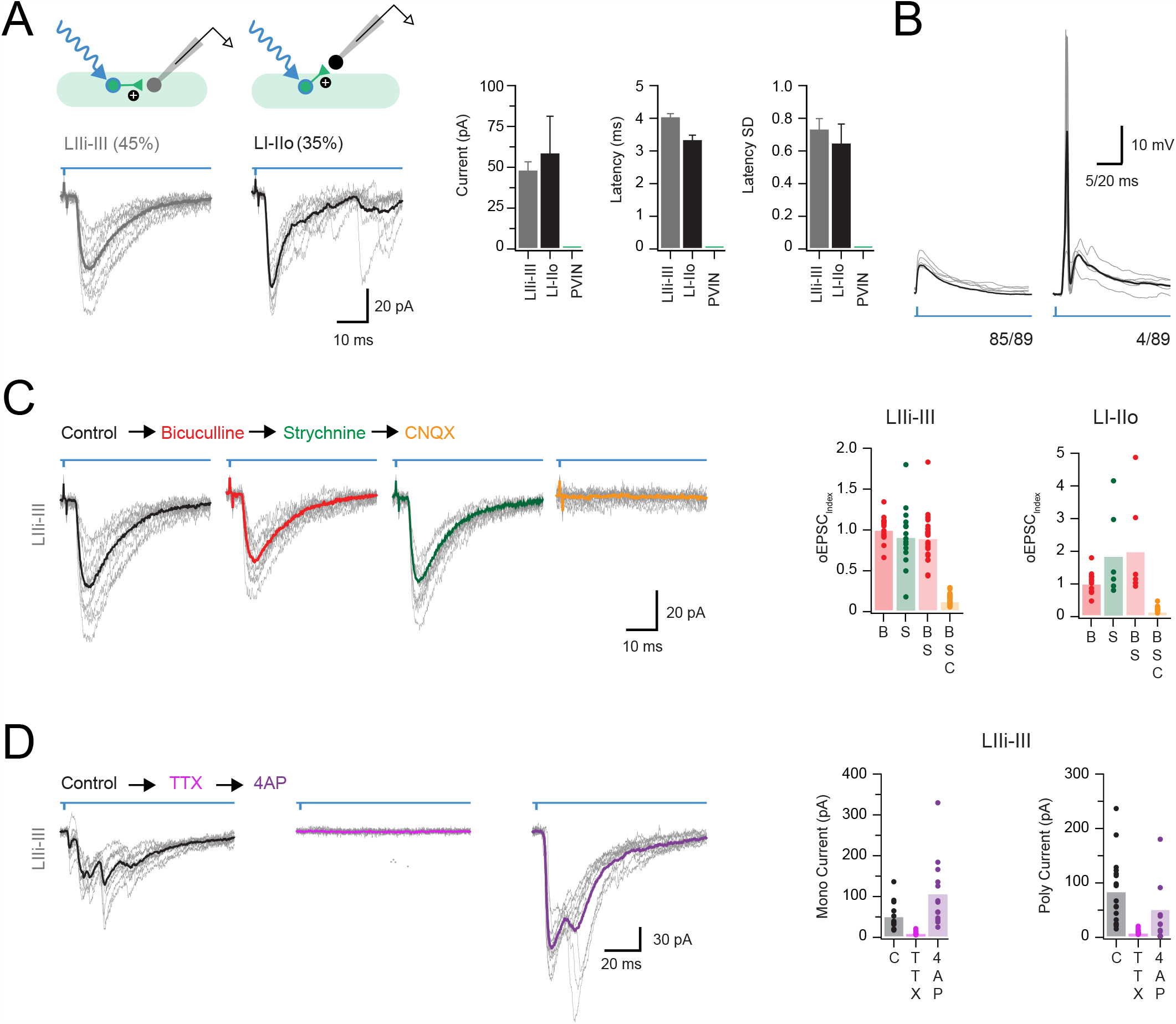
ePVINs are a source of monosynaptic glutamatergic excitation. **A**, Upper schematics show the two recording configurations used to study monosynaptic excitatory connections: PVIN to lamina IIi-III neurons, and PVIN to lamina I-IIo neurons. Voltage clamp recordings (−70 mV) showing optically-evoked excitatory postsynaptic currents (oEPSCs) in neurons from laminae I-IIo and IIi-III, but not in PVINs as no oEPSC responses were observed in this population (10 consecutive sweeps and average overlayed). Group data (right) compares oEPSC amplitude, latency, and latency standard deviation (jitter), highlighting oEPSC monosynaptic characteristics. **B**, Current clamp recordings (−60 mV) showing photostimulation-evoked oEPSCs rarely induce action potential discharge in postsynaptic neurons (only 4/89 recordings featured AP discharge in oEPSP responses). **C**, Representative oEPSCs recorded from a single neuron (black), and following sequential addition of bicuculline (red), strychnine (green), and CNQX (orange). Block of GABA and glycine receptors has minimal effect on oEPSCs, while addition of CNQX abolishes the response. Group data (right) shows effects of this drug application approach in neurons from laminae IIi-III (left) and I-IIo (right). **D**, Representative oEPSCs recorded from a single neuron (black) and following sequential addition of TTX (pink) and 4AP (purple). Group data below shows effects of drug application on monosynaptic current (bottom left) and polysynaptic current (bottom right). Addition of 4AP following TTX application recovers monosynaptic oEPSCs, as well as many presumably action potential-independent polysynaptic oEPSCs.

### PVIN evoked polysynaptic input arises from multiple distinct circuits

In addition to short latency (monosynaptic) oEPSCs identified above, PVIN photoactivation evoked polysynaptic excitatory input (oEPSCs) in 38% (87/231) of UN:LIIi-III cells, 49% (31/63) of UN:LI-IIo, and 50% (18/36) of PVINs (Figure 7A). These oEPSC had longer latencies (16.9 ± 1.6 ms, 19.5 ± 2.2 ms, and 8.2 ± 0.6 ms, UN:LIIi-III, UN:LI-IIo, and PVINs respectively) and exhibited greater jitter (4.3 ± 0.71 ms, 9.3 ± 1.6 ms, and 0.98 ± 0.27 ms) than monosynaptic oEPSCs. Polysynaptic oEPSC amplitude varied across the sample, with PVINs receiving the largest input followed by UN:LIIi-II recordings and finally UN:LI-IIo cells (235 ± 87 pA *vs*. 140 ± 52 pA *vs*. 68 ± 15.5 pA, respectively, p = 0.15), however, all responses were abolished by bath applied CNXQ (oEPSC_index_: 0.11 ± 0.03 p < 0.001, 0.16 0.04 p < 0.001, and 0 ± 0, p < 0.001; Figure 7B-D). Such responses clearly required photostimulation to recruit glutamateric circuitry, with the ePVINs described above one obvious source of input. Despite this, monosynaptic oEPSCs rarely initiated AP discharge (Figure 6B) arguing against ePVINs being a major contributor to the polysynaptic oEPSC responses observed.

**Figure 7.**
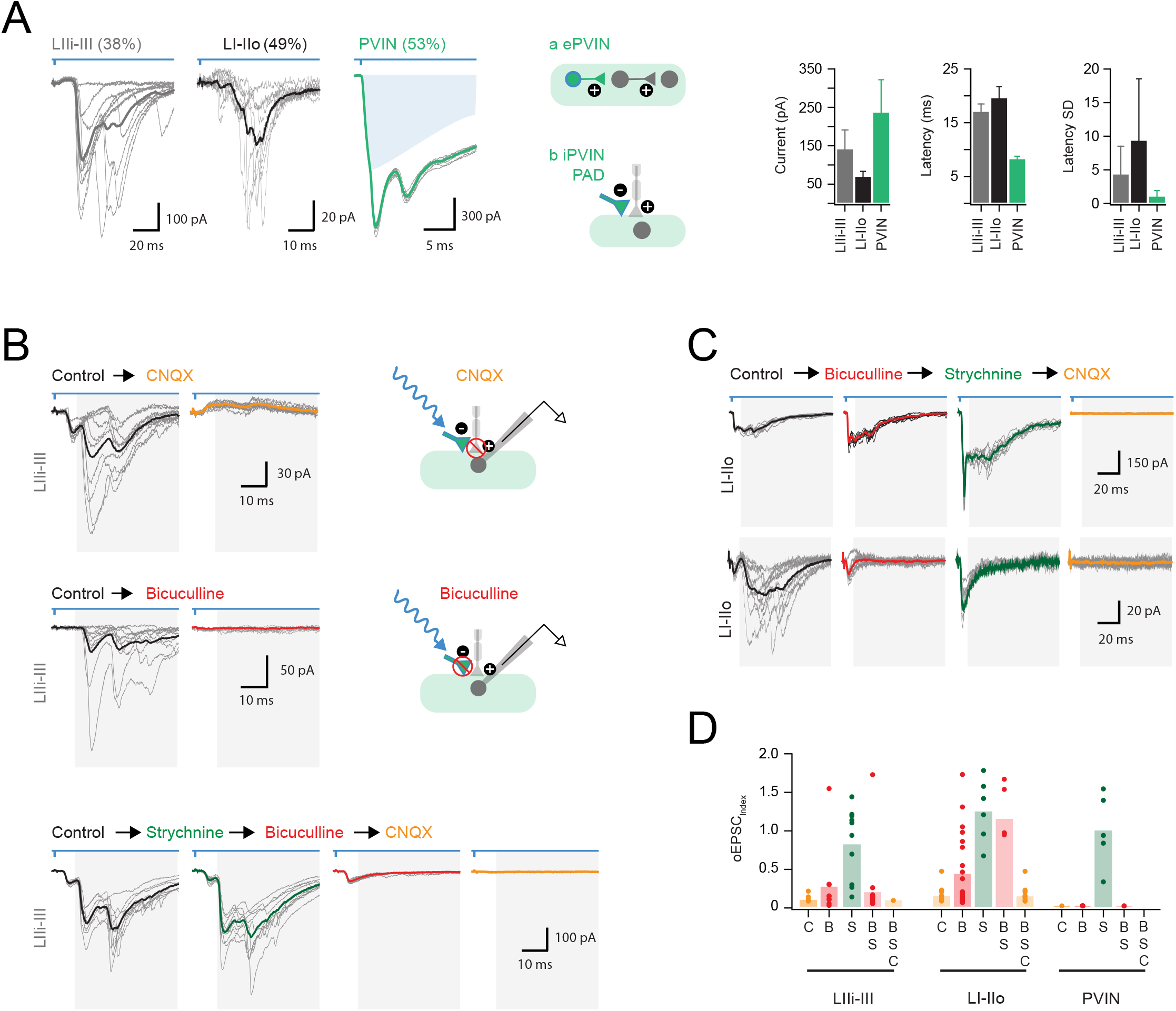
PVIN activation evokes polysynaptic EPSCs. **A**, Voltage clamp recordings (−70 mV) showing optically-evoked excitatory postsynaptic currents (oEPSCs) in neurons from laminae I-IIo and IIi-III, as well as PVINs (10 consecutive sweeps and average overlayed). Note these excitatory responses exhibited long latencies. Blue shading in PVIN trace denotes underlying photocurrent isolated by pharmacological block of synaptic events. Schematics (middle) shows potential polysynaptic mechanism for these oEPSCs. Responses could arise from photostimulation of and excitatory PVIN that recruits an interposed excitatory IN producing the recorded response (**a**, middle upper); or photostimulation may activate an inhibitory PVIN input onto a primary afferent fibre, that in turn on the recorded neuron (**b**, middle lower). Group data (right) compares oEPSC amplitude, latency, and latency standard deviation (jitter). These characteristics are consistent with polysynaptic oEPSCs. **B**, Representative oEPSCs recorded from neurons in laminae IIi-III (black traces) following addition of CNQX (upper left, orange trace) and bicuculline (upper right, red trace), with schematics (right) showing the postulated circuit and site of drug action. Sequential addition of strychnine (lower, green), bicuculline (lower, red), and CNQX (lower, orange). GABA and AMPA receptor block abolishes polysynaptic oEPSCs, while addition of strychnine has minimal effect. **C**, Representative oEPSCs recorded from neurons in lamina I-IIo (black), with sequential bath application of bicuculline (red), strychnine (green), and CNQX (orange). Two series of traces highlight the variability in drug responsiveness of neurons in laminae I-IIo. Addition of bicuculline had minimal effect in some recordings (upper traces), whereas bicuculline abolished polysynaptic oEPSCs in many recordings (lower traces). **D**, Group data shows effects of drug application in neurons from laminae IIi-III (left), I-IIo (middle), and PVINs (right). Note CNQX or bicuculline abolished all polysynaptic responses in PVINs and most neurons in IIi-III, but responses to bicuculline varied in recordings from lamina I-IIo.

Presynaptic inhibition arising from iPVINs is another mechanism that could contribute to these polysynaptic excitatory responses under our experimental conditions (Fink et al., 2014; Boyle et al., 2019). Specifically, iPVIN photostimulation has been shown to evoke GABA-mediated primary afferent depolarisation (PAD) that is capable of eliciting neurotransmitter release from afferent terminals (Boyle et al., 2019). This mechanism would produce longer latency polysynaptic oEPSCs as we have previously demonstrated in iPVIN photostimulation responses recorded in lamina II vertical neurons (Boyle et al., 2019). These previous experiments showed that such connectivity is sensitive to glutamatergic, as well as GABA_A_ receptor blockade at the inhibitory PVIN-to-primary afferent synapses. Thus, we assessed the sensitivity of polysynaptic oEPSCs recorded here to bath applied bicuculline (Figure 7B-D). All polysynaptic oEPSCs in PVINs were abolished by bicuculline (oEPSC_index_: 0 ± 0, p < 0.001), which also significantly reduced polysynaptic oEPSCs in UN:LIIi-III and UN:LI-IIo cells (oEPSC_index_: 0.28 ± 0.16, p = 0.002 and 0.45 ± 0.11, p < 0.001, respectively). These data support a role for iPVIN-evoked PAD events in generating polysynaptic oEPSCs. Furthermore, given the axoaxonic nature of iPVIN input to primary afferents, it is possible that polysynaptic oEPSCs arising from these connections may persist in the presence of TTX and 4-AP. Indeed, polysynaptic oEPSCs were still detected in the presence of TTX and 4-AP in some recordings (8/18). This result supports the interpretation that the polysynaptic circuitry in question does not rely on spiking in an interposed neuron, but rather PAD in primary afferent terminals (Figure 6D). Taken together, these findings confirm iPVIN activation during photostimulation causes PAD, and that this event is capable of producing polysynaptic oEPSCs by driving neurotransmitter release from primary afferent terminals under these experimental conditions.

The above pharmacology is particularly interesting because, although a substantial literature suggests presynaptic inhibition is mediated by GABA, our experiments on postsynaptic inhibition identified glycine as the dominant neurotransmitter used by PVINs (Figure 5B-C). Thus, the impact of glycine receptor block on polysynaptic oEPSCs was also assessed (Figure 7B-D). Unlike bicuculline, strychnine did not affect polysynaptic oEPSC amplitude in UN:LIIi-III, UN:LI-IIo, and PVIN recordings (EPSC_index_: 0.83 ± 0.15, p = 0.28, 1.26 ± 0.17 p = 0.192, and 1.00 ± 0.22, p = 1.0, respectively). This is consistent with GABA being the sole mediator of iPVIN-mediated presynaptic inhibition, however, it also is worth noting that the EPSC_index_ increased following administration of strychnine in some UN:LIIi-III (5/10 cells tested), UN:LI-IIo (4/6 cells tested), and PVINs (2/5 cells tested) (Figure 7D). These data suggest that glycinergic inhibition also regulates this circuitry, likely via ongoing tonic and phasic glycinergic inhibition of the photostimulated PVINs (Gradwell et al., 2017).

Despite the above evidence for PAD-evoked polysynaptic oEPSCs, addition of bicuculline did not block all polysynaptic oEPSCs recorded from either UN:LIIi-III (5/11) or UN:LI-IIo (8/11), but did in PVINs (9/9) (Figure7C-D). Thus, our data also provides evidence that photostimulation of ePVINs produces some signalling through polysynaptic circuits. On some occasions, disinhibition produced by bicuculline and strychnine even unmasked polysynaptic excitatory responses in UN:LIIi-III (2/11) and UN:LI-IIo cells (3/11), suggesting existing excitatory pathways were supressed by ongoing inhibition. It is likely this ongoing inhibition is largely glycinergic, as bicuculline occasionally resulted in complete polysynaptic block only for subsequent strychnine application to unmask polysynaptic excitatory circuit responses. Together these observations indicate that polysynaptic excitatory circuits can also be driven by ePVINs. The output of these excitatory circuits were more likely to be observed in laminae I-IIo, were not observed in PVINs, and can be better resolved under disinhibited conditions.

Complex pharmacology made differentiation of iPVIN and ePVIN evoked polysynaptic oEPSCs difficult to dissect beyond the conclusion that our dataset likely contains polysynaptic oEPSCs originating from both circuits. Despite this, in recordings from UN:LIIi-III cells, oEPSCs driven by ePVIN polysynaptic circuits (bicuculline-insensitive) had longer latency and greater jitter when compared to oEPSCs driven by bicuculline-sensitive iPVIN PAD-mediated polysynaptic circuits (latency: 26.8 ± 6.4 ms vs 12.2 ± 0.8 ms, p < 0.001; jitter: 8.4 ± 3.5 ms vs. 1.8 ± 0.3 ms, p = 0.001). This distinction was less clear in UN:LI-IIo recordings with similar latency (19.7 ± 2.9 ms *vs*. 18.2 ± 2.2 ms) and jitter (9.5 ± 2.1 ms *vs*. 9.3 ± 2.1 ms) for bicuculline-resistant and bicuculline-sensitive polysynaptic oEPSCs observed. The distinction between polysynaptic oEPSC latencies in LIIi-III but not LI-IIo may relate to the termination site of A-LTMRs under iPVIN-mediated PAD control. Specifically, A-LTMRs terminate in deeper lamina and therefore neurons in that region (LIIi-III) more likely to receive direct input from A-LTMRs, resulting in shorter latency iPVIN-mediated polysynaptic oEPSCs, compared to those arising from ePVIN circuits. In contrast, iPVIN-mediated polysynaptic oEPSCs in LI-IIo, where A-LTMR input is rare, likely require recruitment of interposed excitatory interneurons in LIIi-III and subsequent longer latency relay of excitation in LI-IIo circuits. In line with this proposed connectivity, our previous work has demonstrated vertical cells in LII represent such an excitatory population that receives direct afferent input regulated by iPVIN presynaptic inhibition (Boyle et al., 2019).

### Postsynaptic targets of PVIN circuits

Given the substantial heterogeneity of dorsal horn neurons (Graham et al., 2007; Todd 2010) we next assessed the AP discharge patterns of neurons receiving oEPSCs (n = 203, Figure 8) as AP discharge has been used to infer inhibitory or excitatory neuron phenotype (Yasaka et al., 2010; Hughes et al., 2012; Smith et al., 2015). Specifically, tonic firing (TF) and initial bursting (or adaptive firing) discharge is common in inhibitory neurons, whereas delayed firing (DF) is typical of excitatory cells. PVIN photostimulation responses were identified in neurons with a range of AP discharge profiles that included tonic firing (TF), initial bursting (IB), delayed firing (DF), and single spiking (SS). We also identified photostimulation responses in neurons with a distinct AP discharge pattern not commonly described. In these cells a rapid depolarising hump was associated with burst of APs at the beginning of depolarisation (Figure 8A). Given our sampling included recordings form lamina III, these neurons may correspond to phasic cells that exhibit pronounced spike frequency adaptation, reported in the deep dorsal horn (Schneider 2003). Morphological recovery of these neurons, herein termed rapidly adapting (RA), confirmed they exhibited extensive rostrocaudally-oriented dendritic arbours as commonly observed in lamina II islet cells. RA neurons more often received monosynaptic oEPSCs than neurons with other discharge patterns (RA = 67%, 18/27; TF = 30% 19/63; IB = 31% 8/26; DF= 40%, 30/75; SS = 50%, 6/12), though all oEPSCs were similar amplitude (RA = 42 ± 8 pA, TF = 37 ± 7 pA, IB = 27 ± 12 pA, DF = 52 ± 16 pA, and SS = 39 ± 16 pA) (Figure 8B). Thus, ePVINs provide appear to provide similar levels of input to other dorsal horn neurons with both excitatory and inhibitory characteristics.

**Figure 8.**
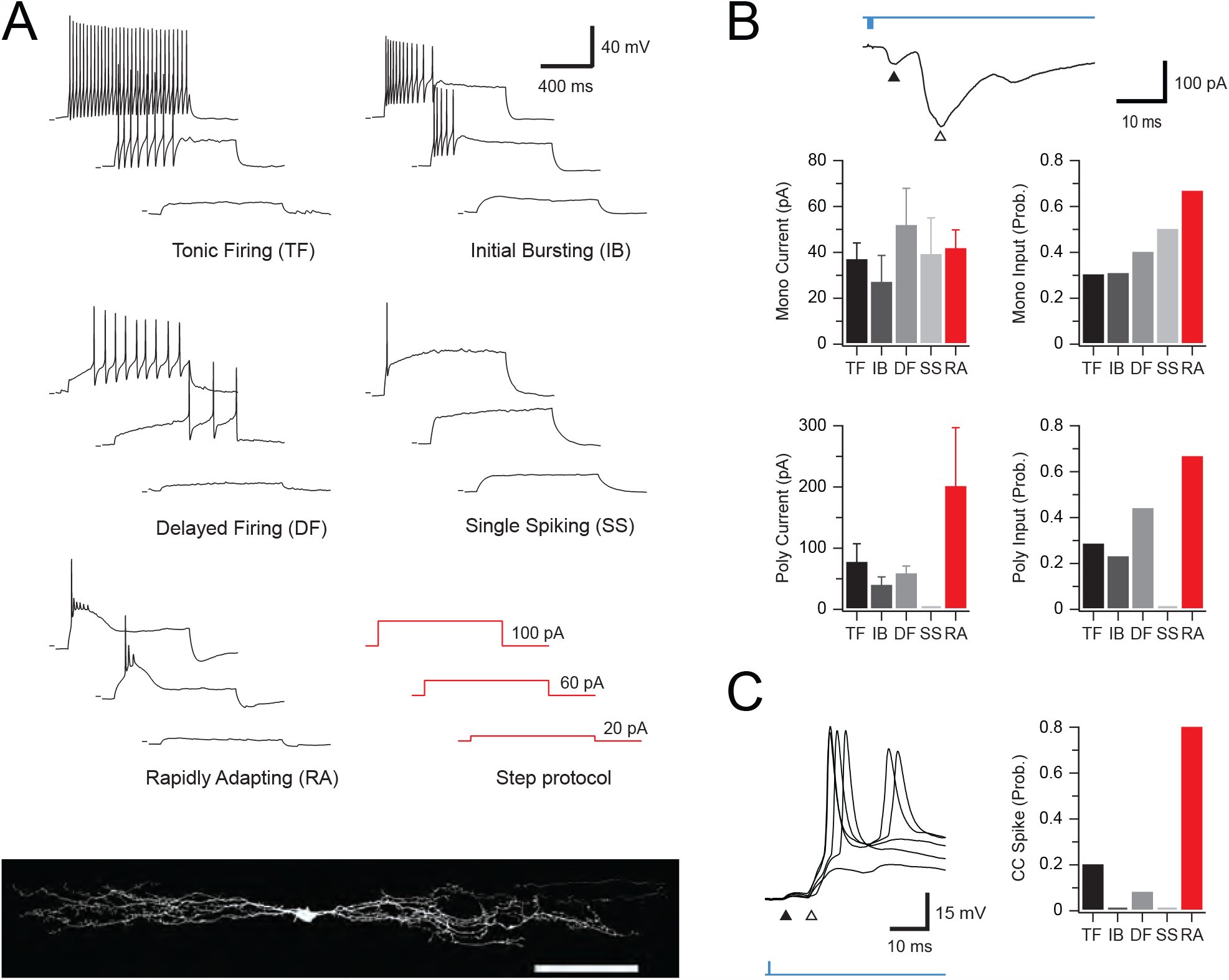
Action potential discharge responses of neurons receiving PVIN mediated glutamatergic inputs. **A**, Example traces of the four characteristic action potential discharge types following depolarizing current injection (bottom right): tonic firing, initial bursting, delayed firing, and single spiking, as well as another, previously unidentified, population of rapidly adapting neurons. Rapidly adapting neurons all exhibited islet cell morphology (bottom), suggesting an inhibitory phenotype. **B**, Top trace is an example voltage-clamp recording from a neuron receiving monosynaptic (black arrow) as well as polysynaptic (white arrow) oEPSC input. Group data graphs (below) show the incidence of monosynaptic (top right) and polysynaptic (bottom right) oEPSCs in neurons with each type of discharge response. This data highlights the higher incidence of polysynaptic oEPSCs in the rapidly adapting population. Similarly, the amplitude of monosynaptic (top left), polysynaptic (bottom left) oEPSCs is compared for responses recorded in each AP discharge category. Only the amplitude of polysynaptic oEPSCs differed where rapidly adapting neurons received larger amplitude inputs. **C**, Trace is an example current-clamp recording from a rapidly adapting neuron receiving monosynaptic (black arrow) as well as polysynaptic (white arrow) oEPSC input. Note the polysynaptic component of the oEPSC reaches AP threshold and evokes a spike. Group data (right) highlights the increased incidence of oEPSC-evoked AP discharge in the rapidly adapting population, versus all other discharge categories.

The AP discharge properties of neurons receiving polysynaptic oEPSCs were also assessed to determine if these connections preferentially targeted excitatory or inhibitory neurons (Figure 8B). As for monosynaptic inputs, RA neurons more often received polysynaptic oEPSCs than neurons with other discharge patterns (RA = 67%, 18/27; TF = 29%, 18/63; IB = 23%, 6/26; DF = 44%, 33/75; SS = 0%, 0/12). The amplitude of polysynaptic oEPSCs was also substantially larger in recordings from the RA population (RA = 201 ± 96 pA, TF = 77 ± 30 pA, IB = 40 ± 14 pA, DF = 59 ± 13 pA), and these oEPSCs were more likely to initiate AP discharge in the RA recordings (RA = 80% 12/15, TF = 20%, 3/15; IB = 0%, 0/4; DF = 8%, 2/25, Figure 8C). In agreement with the postulated role of PVIN-mediated PAD in initiating these polysynaptic oEPSCs, they could be abolished by bicuculine (EPSC_index_: 0.13 ± 0.02, n = 5) or CNQX (EPSC_index_: 0.07 ± 0.01, n = 2), but were unaffected by strychnine (EPSC_index_: 1.15 ± 0.15, n = 3). Thus, RA neurons appear to receive large afferent inputs that are strongly regulated by iPVIN, whereas the afferent input regulated by this mechanism in other inhibitory populations (TF and IB) and excitatory populations (DF) is less pronounced.

### PVIN-mediated input to Lamina I Projection neurons

In light of the diverse dorsal horn circuits regulated by PVINs, we also sought to examine PVIN-mediated input specifically to lamina I projection neurons (PNs). PV^Cre^;Ai32 animals (n=8) received bilateral intracranial virus injections of AAV9-CB7-Cl∼mCherry in the parabrachial nuclei to maximally label PNs and spinal cord slices (both transverse and sagittal planes) were subsequently prepared 2-4 weeks later. Targeted recordings from mCherry-labelled PNs (Figure 9A) used a KGluc-based internal solution with oEPSCs and oIPSCs presenting as inward and outward currents, respectively (holding potentials of −70 mV and −30 mV). Under these conditions, photostimulation of PVINs evoked a range of responses (Figure 9B-C). In transverse slices, monosynaptic oEPSCs exhibited bicuculline-insensitivity (4/33, 12%), whereas bicuculline/strychnine sensitive monosynaptic oIPSCs were never observed (0/28, 0%). In sagittal slices, monosynaptic bicuculline-insensitive oEPSCs (4/11; 36%) and bicuculline/strychnine sensitive oIPSCs (2/8; 25%) were observed in PNs. These pharmacologically identified monosynaptic oEPSCs and oIPSCs exhibited short latencies (3.16 ± 0.38 ms and 3.11 ± 1.10 ms) and limited jitter (0.88 ± 0.27 ms and 0.54 ± 0.08 ms), similar to UN:LIIi-III and UN:LI-IIo cells. Thus, both the ePVINs and iPVINs provide input to PNs in lamina I.

**Figure 9.**
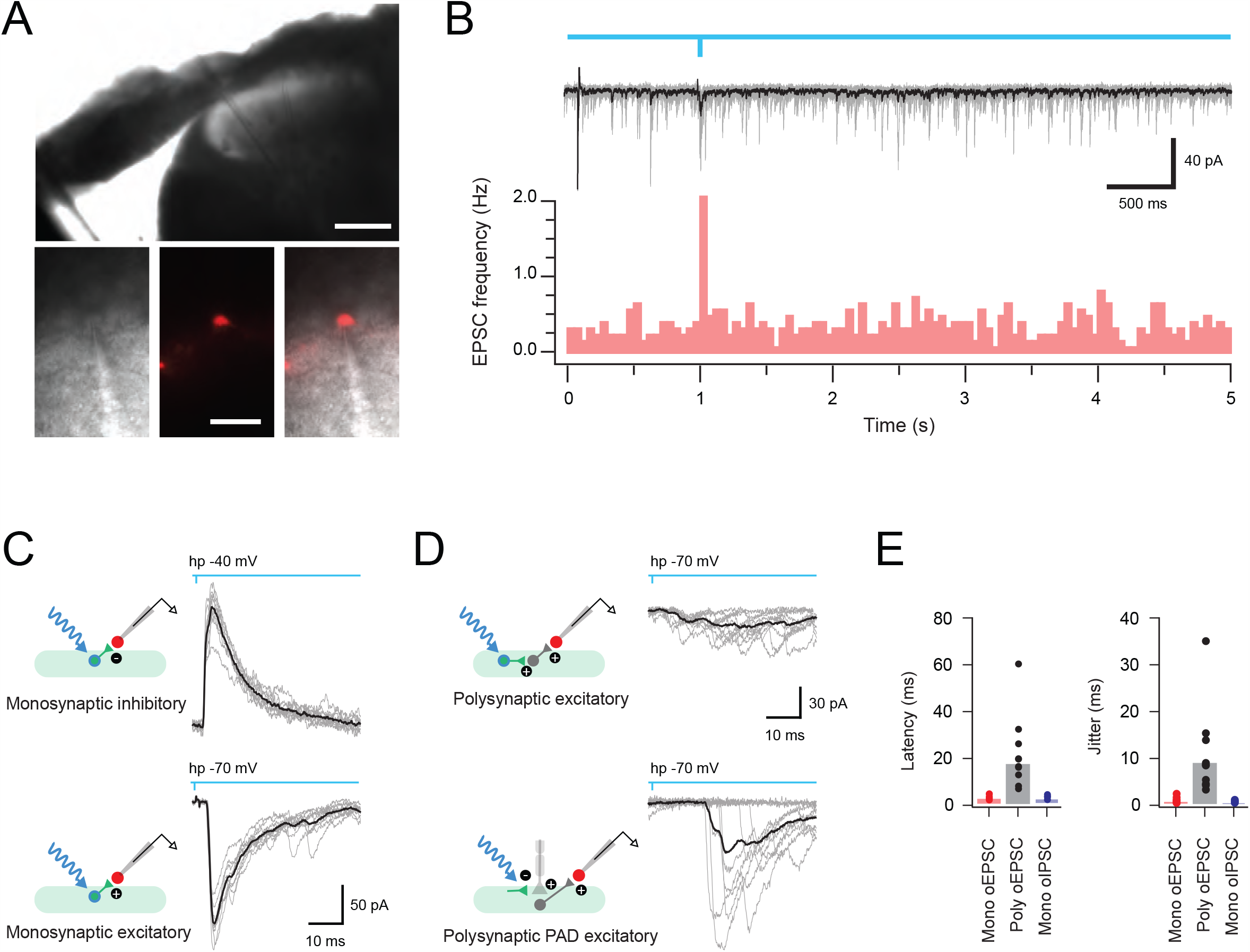
PVIN-mediated input to lamina I projection neurons. **A**, low magnification image (upper) shows transverse spinal cord slice with recording pipette in place to record from a lamina I projection neuron (PN), with higher magnification images showing the targeted recording configuration between the recording electrode and retrogradely-labelled PN (brightfield left, mCherry fluorescence centre, overlay right). **B**, 10 overlayed traces with superimposed average (grey and black, upper) along with peristimulus histogram from this data (below) show ongoing EPSC frequency recorded from a PN with brief (1 ms) photostimulation of PVINs applied one second into the trace (blue line). **C**, Photostimulation responses recorded in PNs showing characteristics of monosynaptic inhibitory (upper) and excitatory (lower) connections. Overlayed traces (grey) show 10 photostimulation (blue) trials on an expanded times scale with averaged responses (black) superimposed. Schematics (left) summarise the postulated underlying circuits between PVINs (green) and PNs (red). **D**, Photostimulation responses recorded in PNs showing characteristics of polysynaptic connections. Overlayed traces (grey) show 10 photostimulation (blue) trials on an expanded times scale with averaged responses (black) superimposed. Schematics (left) summarise the postulated underlying circuits between PVINs (green) and PNs (red). Polysynaptic responses could be further differentiated into those that arose from excitatory circuits evoked by photostimulation of ePVINs (upper), and those that arose from photostimulation of iPVINs causing primary afferent depolarisation (PAD) and subsequent excitatory signalling from these terminals (lower). **E**, Group data plots compare latency and jitter of photostimulation responses in monosynaptic excitatory, polysynaptic excitatory and monosynaptic inhibitory connections. Scale bars in A (μm) = 100 upper; 40 lower.

In addition to monosynaptic inputs, some PN recordings (10/44; 22%) exhibited polysynaptic oEPSCs (Figure 9D-E) based on a longer latency (17.81 ± 3.56 ms) and enhanced jitter (9.14 ± 1.53 ms). As above, polysynaptic oEPSCs may arise from distinct circuits involving ePVINs or iPVIN-mediated PAD, differentiated by their bicuculline sensitivity. Bicuculline reduced or abolished polysynaptic oEPSC amplitude in 3/6 PNs tested, and subsequent application of CNQX abolished remaining polysynaptic oEPSCs. Bicuculline sensitivity indicates a polysynaptic circuit through iPVIN-mediated PAD of primary afferents that either directly terminates on PNs or provides input via an interposed excitatory interneuron (Figure 9D, lower). As noted above, our recent work has identified vertical cells, an excitatory interneuron population with axons arborising in lamina I, as a likely candidate to complete such a circuit (Boyle et al., 2019). Finally, the presence of bicuculline-insensitive oEPSCs (3/6) implies that polysynaptic input to PNs is also mediated by ePVINs (Figure 9D, upper).

### PVINs regulate nociceptive circuits

The above results confirm a significant fraction of unidentified cells in laminae I-IIo, and of PNs, received mono- and polysynaptic input from both ePVINs and iPVINs. Given many cells in this region receive nociceptive input, this suggests PVINs play a role in nociceptive processing as well as their established role in gating innocuous tactile inputs. To test this, a subset of recordings assessed the effect of bath-applied capsaicin on miniature excitatory postsynaptic current (mEPSC) frequency in UN:LI-IIo cells that received PVIN-mediated oEPSCs or oIPSCs (Figure 10, n = 9). Under these conditions capsaicin causes selective increase to mEPSC frequency only in neurons that receive direct input from TRPV1+ (nociceptive) inputs. Bath-applied capsaicin increased mEPSC frequency in these recordings (4.28 ± 2.31 *vs*. 10.33 ± 4.84 Hz, p = 0.049) without altering amplitude (15.4 ± 1.2 *vs*. 17.2 ± 1.9 pA, p = 0.164) or time course (rise time = 1.27 ± 0.14 ms *vs*. 1.20 ± 0.15 ms; decay time constant = 4.54 ± 0.48 ms *vs*. 4.03 ± 0.45 ms), confirming the sample included neurons with nociceptive input (Figure 10A). As mEPSC frequency fluctuates and the effect of capsaicin varied, a threshold was set (mEPSC frequency increase of 3 standard deviations above the mean baseline rate) for considering a neuron as receiving capsaicin-sensitive input. Using this criterion two thirds of UN:LI-IIo cells (6/9) received capsaicin-sensitive input. Assessment of PVIN photostimulation in these recording showed 1/6 received monosynaptic iPVIN-mediated oIPSCs (Figure 10B), 4/6 received monosynaptic ePVIN-mediated oEPSCs (Figure 10C), and 2/6 received bicuculline-sensitive polysynaptic oEPSCs (Figure 10D). Taken together, these data suggest roles for both ePVINs and iPVINs modulating nociceptive circuits.

**Figure 10.**
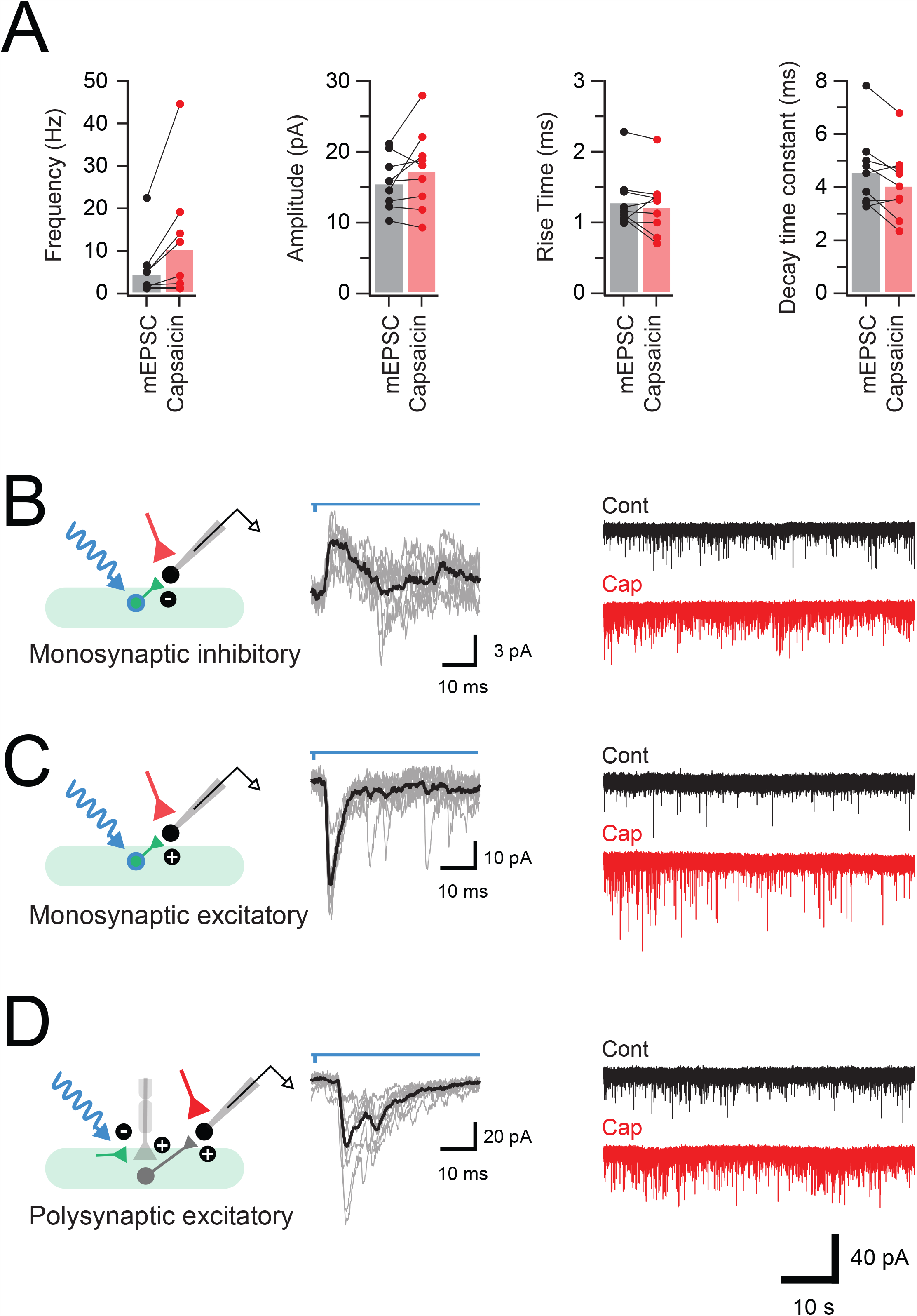
PVINs modulate nociceptive circuits. **A**, plots show group data, demonstrating the increase in mEPSC frequency following bath applied capsaicin. Neurons were deemed to be capsaicin sensitive if capsaicin application increased mEPSC frequency by three standard deviations or more above the mean baseline rate. mEPSC amplitude and rise time remained unchanged, whereas decay time constant was reduced following capsaicin application. **B-C**, Schematics (left) summarise the microcircuits producing photostimulation-evoked postsynaptic currents (oPSCs, 10 consecutive sweeps and average overlayed) in neurons from lamina I-IIo (middle). Continuous traces (right) show mEPSC recordings (TTX 1 μM, bicuculline 10 μM, and strychnine 1 μM) from corresponding neurons before (black) and after (red) bath application of capsaicin (2 μM). Capsaicin application increases mEPSC frequency without altering amplitude, confirming these neurons received nociceptive input. These recordings identified some neurons that received monosynaptic oIPSCs (**B**, short latency outward currents during voltage clamp at −40 mV) presumably arising from photostimulation of inhibitory iPVINs; neurons that received monosynaptic oEPSCs (**C**, short latency inward currents during voltage clamp at −70mV) arising from photostimulation of an ePVIN population; or neurons that received polysynaptic oEPSCs (**D**, longer latency inward currents during voltage clamp at −70mV) arising from photostimulation of iPVIN terminals cause primary afferent depolarization (PAD) and synaptic transmission at those terminals.

### PVIN activation suppresses AP discharge in other DH populations

Given the complex range of signals produced by PVIN phototstimulation, the combined impact of activating this circuitry on DH neuron discharge was assessed. In separate protocols, UN:LIIi-III cells received a series of long (Figure 11A, n=9) or short (Figure 11B, n=14) depolarising current step injections to evoke AP discharge (1 s and 50 ms duration, respectively; 20 pA increments). These steps were repeated three times with PVIN photostimulation delivered immediately prior to the second (middle) step series. PVIN photostimulation prior to long step depolarisations significantly increased the latency of AP discharge (72.9 ± 16.4 ms *vs*. 134.3 ± 27.9, p = 0.013 *vs*. 76.8 ± 17.6 ms, p = 0.017; step 1, step 2, and step 3, respectively), but did not affect rheobase current (100 ± 28 pA *vs*. 100 ± 28 pA, p =1.0 *vs*.100 ± 28 pA, p =1.0), or the number of APs evoked per step (3.7. ± 1.2 *vs*. 3.3 ± 0.9, p = 0.397 *vs*. 3.1 ± 0.5, p = 0.746). In short step responses, prior PVIN photostimulation increased rheobase current (89 ± 11 pA *vs*. 130 ± 17 pA, p = 0.012 *vs*. 87 ± 10 pA, p = 0.011 for step 1, step 2, and step 3), increased the latency to AP discharge (24.0 ± 3.4 ms *vs*. 39.5 ± 3.7 ms, p = 0.001 *vs*. 22.6 ± 2.6 ms, p < 0.001), and reduced the number of APs in the response (2.1 ± 0.2 *vs*. 1.3 ± 0.2, p = 0.003 *vs*. 1.9 ± 0.2, p = 0.045). Thus, activation of PVINs produced an overall inhibition of AP discharge in unidentified circuits within laminae IIi-III.

**Figure 11.**
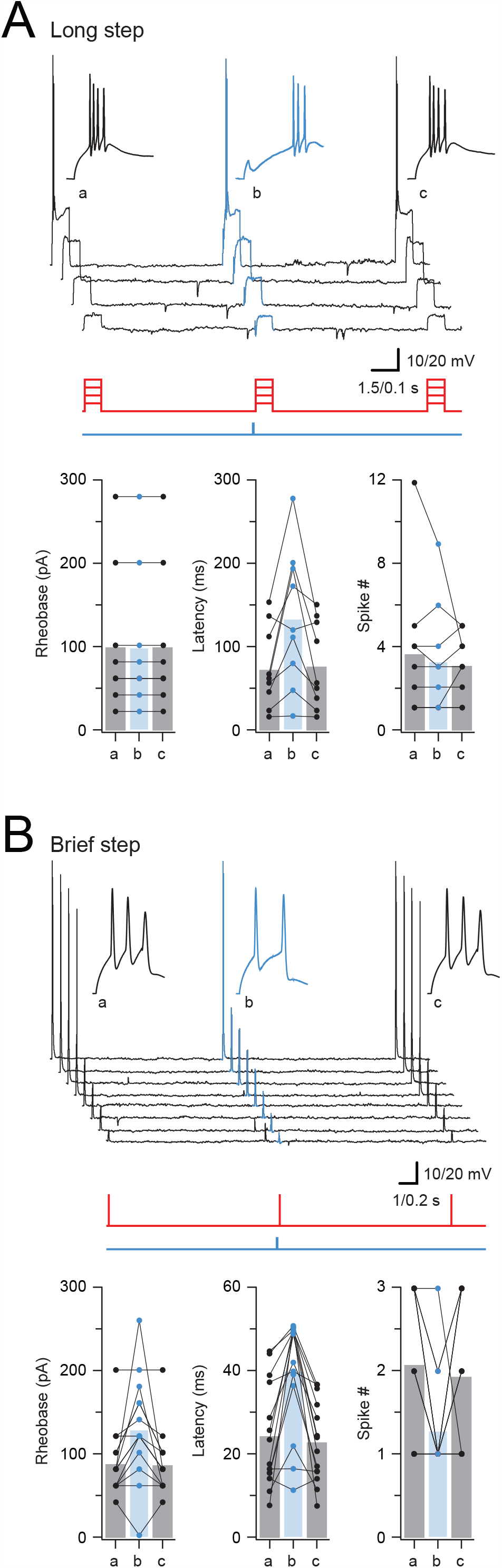
PVIN-mediated inhibition alters AP discharge. **A**, top trace shows AP discharge responses recorded in a neuron from lamina IIi-III during long (1s) depolarizing step injections: (a) pre-test step; (b) test step – preceded by a single PVIN photostimulation (10 ms duration at the onset of depolarizing current steps); and (c) post-test step. Traces below show current step injection (red) and photostimulation timing (blue). Insets show the onset of AP discharge on expanded time scale highlighting the delay to AP discharge in the test response (b). Plots below show group data comparing rheobase current, AP latency and AP number between the pre-test, test and post-test step responses. Only spike latency differed between conditions. **B**, shows same general experimental approach as in A, however depolarizing current injections were short duration (50ms). This paradigm highlighted more significant modulation of AP discharge. Specifically, in response to shorter depolarising step (more reflective of the time course likely in synaptic excitation), rheobase, latency and number of spikes in response to stimulation are all altered by preceding activation of PVINs and the associated inhibition they mediate. Together, responses show PVIN-mediated inhibition is capable of modulating the timing of AP discharge following prolonged depolarization, as well as the rheobase for AP discharge and number of APs produced following brief depolarization.

### Circuit mapping PVIN targets in vivo

We assessed the pattern of Fos labelling following *in vivo* photostimulation PVINs in anaesthetised PV^Cre^;Ai32 mice (n = 3) to determine the pattern of spinal activation following selective recruitment of these cells. The dorsal spinal cord (L4 segment) of these animals was exposed by laminectomy and an optic fibre (400μm diameter) applied unilateral photostimulation (10 ms pulses @ 10 Hz for 10 minutes) as described previously (Smith et al., 2019). Animals were maintained under anaesthesia for a further 2 hours before overdose and transcardial perfusion fixation. Subsequent tissue processing for Fos protein expression mapped photostimulation induced activation within the dorsal horn. This protocol produced reliable Fos expression, discretely restricted to the area underlying the optic fibre (Figure 12AB). Most Fos positive profiles were located in lamina I-IIo outside the PVIN plexus, with some occasional immunolabelled cells also detected in the deeper dorsal horn, laminae III and IV. Surprisingly, no Fos labelling was detected in ChR2-YFP expressing PVINs. The absence of Fos expression in YFP-expressing cells was unexpected but is consistent with the extensive connectivity both between iPVINs and with ePVINs (Figure 3) and the dominance of PVIN-mediated inhibition in this region (Figure 12C-D). The release of GABA and glycine from axon terminals of iPVINs and the resultant widespread inhibition is likely to prevent their postsynaptic targets from reaching the activity threshold required for Fos expression. In contrast, Fos-expression in dorsal horn neurons that did not express YFP (channelrhodopsin-2) was expected and could result from glutamatergic (excitatory) inputs derived from two principal candidates. The most likely source would be excitatory inputs derived from axon terminals of ePVINs. Photostimlation can trigger transmitter release from these axon terminals, even in the absence of action potentials, and this would result in activation of their postsynaptic targets. A second source of excitatory input in this experimental preparation may arise from myelinated primary afferents under presynaptic control from iPVINs. Under appropriate experimental conditions, photostimlation of axoaxonic synapses induces the release of glutamate from primary afferent terminals (Fink et al., 2014; Boyle et al. 2019). While we cannot exclude a role for this iPVIN-mediated excitatory signal in our *in vivo* preparation, previous work has shown this presynaptic-inhibition mediated signal to be temperature sensitive and diminished/abolished at physiological temperature. This observation leaves photostimulation of ePVINs the most likely source of Fos induction.

**Figure 12.**
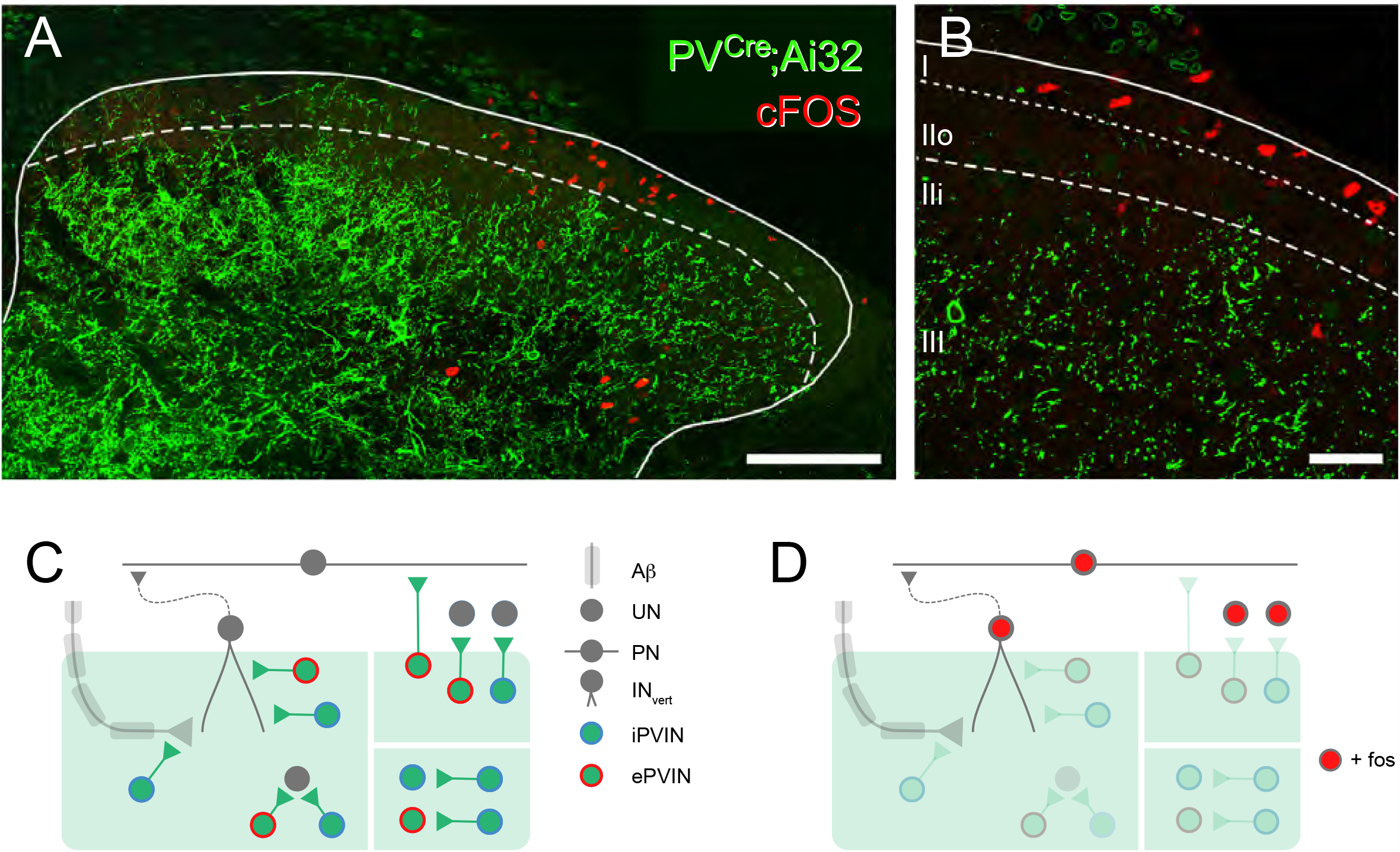
Spinal activation patterns following in vivo photostimulation of PVINs. **A**, image shows maximum intensity projection of a spinal cord section taken from a PVCre;Ai32 mouse that underwent unilateral *in vivo* spinal photostimulation under anaesthesia. Immunolabelling for the activity marker Fos (red), and GFP to locate PVIN cells and processes (green) shows Fos expression in cells predominantly located in laminae I and IIo, with only scattered Fos positive cells in deeper laminae. Note the absence of Fos expression in the PVIN plexus. **B**, higher magnification single optical section from tissue in **A** with lamina boundaries superimposed shows strong Fos expressing profiles located in lamina I, implicating these cells in nociceptive processing. **C**, schematic summarises identified iPVIN and ePVIN connections. ePVINs and iPVINs provide input to interneurons in LIIi-LIII including vertical cells (Boyle et al., 2019), and iPVINs also provide presynapticinput to myelinated primary afferents (left). iPVINs provide strong inhibition to neighbouring PVINs in LIIi-LIII (lower right). ePVINs and iPVINs provide input to interneurons and projection neurons located in LI-LIIo (upper right). **D**, schematic summarises activation of DH neurons following in vivo photostimulation of PVINs. Circuits inhibited by PVINs in LIIi-LIII and therefore unlikely to reach threshold for Fos expression are faded. In contrast, Fos expression is most pronounced in circuits in LI-LIIo, where ePVIN outputs are more likely to excite cells (including projection neurons). Scale bar (µm): A = 100; B = 20.

## Discussion

In this study, we have used *in situ* hybridisation protocols to establish that parvalbumin-expressing interneurons in the spinal dorsal horn are comprised of two similar-sized populations of excitatory (glutamatergic) and inhibitory (GABAergic) interneurons. These populations can be defined by their morphological properties, with iPVINs having larger soma and more extensive (elongated) dendritic arbors than ePVINs. Detailed anatomical analysis also showed that both ePVINs and iPVINs form homotypic and heterotypic synaptic connections with other PVINs in in laminae IIi and III. We have used optogenetic approaches to show that glutamatergic neurotransmission from ePVINs is distributed throughout the dorsal horn, most prominent in nociceptive circuits and exciting both excitatory interneurons in lamina IIo and lamina I projection neurons that relay information to the parabrachial nucleus. We also determined that inhibition mediated by iPVINs is more prominent in laminae IIi and III, and targets both the central terminals myelinated afferents as well as many dorsal horn interneurons. PVIN-mediated presynaptic inhibition of primary afferent input relies solely on GABAergic signalling, whereas postsynaptic inhibition of dorsal horn interneurons is mediated via both GABA and glycine transmission, with glycinergic signalling predominating. This data highlights substantial diversity and selectivity in the neurotransmitters used by PVINs in dorsal horn circuits and provides important information on how these populations help integrate and modulate somatosensory information.

### Neurotransmitter heterogeneity of PVINs in the mouse dorsal horn

Our finding that half the PVINs in mouse laminae IIi-III are glutamatergic was unexpected given that previous studies have reported the majority of PVINs to be inhibitory. Immunohistochemical studies in the rat show that ∼75% of PVINs in laminae I-III co-express GABA and/or glycine (Antal et al., 1991; Laing et al., 1994), although similar estimates in the mouse have varied considerably. One study reported that ∼95% of PVINs captured in a PV^Cre^; Ai14 cross co-express GABA and glycine, and that less than 5% of PVINs captured in this line show immunolabelling for Lmx1b, a developmental marker of excitatory interneurons (Petitjean et al., 2015). These findings imply that PV-expression is confined largely to inhibitory interneurons, although other lines of evidence have since proposed that glutamatergic PVINs are more abundant. In one study where a PV-tdTomato BAC transgenic mouse line was crossed with either VGLUT2^iresCre^ or VGAT^iresCre^ mice labelled with Cre-dependent reporters, it was estimated that 36% of PVINs are glutamatergic (Abraira et al., 2017). Transcriptomic studies in the mouse dorsal horn have also reported PV expression in both excitatory (Glut1 and Glut2) and inhibitory (Gaba 10, and Gaba13-15) populations, and show that cells within each of these subtypes were common within laminae II and III (Häring et al., 2018). It is likely that discrepancies in the relative incidence of glutamatergic PVINs reflect differences in fidelity of the various transgenic mouse lines used to capture these populations, and this serves to highlight the need for careful validation of genetic labelling patterns with either immunohistochemistry or *in situ* hybridisation. The discrepancies in the literature prompted us to address the question directly using approaches where PVINs were targeted using immunohistochemistry and *in situ* hybridisation. Both these approaches yielded broadly similar results, where ePVINs accounted for approximately half of PVINs in LIIi-III. These findings demonstrate that ePVINs have been underrepresented in classification schemes of dorsal horn interneurons and warrant further investigation to define their roles in dorsal horn circuitry.

### Excitatory and inhibitory PVIN connectivity

Our study provides the first insight into the synaptic connectivity of ePVINs and the circuits they comprise. We show that ePVIN derived excitation onto dorsal horn neurons is distributed throughout the dorsal horn including laminae I and IIo as well as deeper laminae (IIi-III), but oEPSCs were typically of small amplitude and rarely evoked AP discharge in postsynaptic targets. These findings suggest that ePVINs serve an integrative role, where their activity sums with other excitatory signals to influence dorsal horn signalling. Recent work on gastrin-releasing peptide (GRP) expressing excitatory interneurons in the dorsal horn has also highlighted how peptide co-transmission can facilitate postsynaptic discharge (Pagani et al., 2019). Specifically, this work showed that low frequency photostimulation of GRP cells produced subthreshold postsynaptic responses, whereas high frequency photostimulation facilitated release of both glutamate and GRP to potentiate subthreshold glutamatergic input and evoke postsynaptic spiking. A similar mechanism could also enhance postsynaptic responses to ePVIN input given our finding that most PVINs also express CCK (Figure 1). The release of CCK has been implicated in the development of tactile allodynia (Kim et al., 2009), and molecular studies report that several superficial dorsal horn populations (including lamina I projection neurons) express both CCKA and CCKB receptor subtypes (Häring et al., 2018). CCK is distributed widely in the mouse dorsal horn and these cells account for up to a third of excitatory interneurons in lamina III (Gutierrez-Mecinas et al., 2019). Further, a distinct subpopulation of CCK-expressing cells that lack PKCγ have recently been found to play a key role in the development of mechanical allodynia in both inflammatory and neuropathic pain states (Peirs et al., 2020). Given that these cells are found primarily in lamina III, anatomically overlapping with the ePVIN population in laminar distribution and transcriptomic groupings (Group 1 and 2; Häring et al., 2018), it is tempting to speculate that PV expression is another defining feature of these cells.

A surprising difference observed between our Brainbow mapping data and optogenetic experiments was the lack of functional evidence for excitatory inputs between PVINs despite anatomical evidence for these connections (Figure 3). While this is difficult to reconcile, it is possible that the excitatory synapses we find in the anatomical studies represent AMPA-lacking “silent synapses” reported in the rat (Bardoni et al., 1998; Li and Zhuo, 1998; Jung et al., 2005), although the presence of such synapses in laminae I and II of adult rats has been contested (Yasaka et al., 2007). Given that we were able to record monosynaptic excitatory connections between ePVINs and other populations of neurons, it remains to be determined whether the synapses between ePVINs and other PVINs do indeed lack AMPA receptors or if such connections are relatively rare and under-sampled in our recordings. By contrast, direct inhibitory synaptic connections between PVINs were common in our data. These observations were in agreement with our Brainbow mapping experiments and mirror anatomical findings for iPVINs in the dorsal horn (Hughes et al., 2012) and a number of other CNS regions (Tamas et al., 2000; Woodruff and Sah, 2007). Unlike iPVINs in other regions, however, we found no evidence for electrical coupling between PVINs or for autaptic synapses reported for PV-expressing cells in the hippocampus and cortex (Meyer et al., 2002; Pawelzik et al., 2003; Manseau et al., 2010; Deleuze et al., 2014). Nevertheless, an important role for synaptic coupling between PVINs has been put forward in driving rhythmic CNS activity patterns including gamma oscillations in the cerebral cortex (Sohal et al., 2009) as well as theta rhythms and theta/gamma synchronisation in the hippocampus (Freund, 2003; Klausberger and Somogyi, 2008; Wulff et al., 2009). This rhythmic activity is thought to be critical for enhanced information coding, including features of normal sensory processing and cognition (Varela et al., 2001; Womelsdorf et al., 2007; Sohal et al., 2009). Comparable rhythmic activity has been reported in the spinal dorsal horn both *in vivo* (Sandkuhler and Eblen-Zajjur, 1994; Eblen-Zajjur and Sandkuhler, 1997) and *in vitro* (Asghar et al., 2005; Chapman et al., 2009), with proposed roles in modality specific coding and information gating (Ruscheweyh and Sandkuhler, 2005). This is consistent with the idea that iPVINs play a ‘gatekeeping’ role in segregating tactile and noxious signalling (Hughes et al., 2012; Petitjean et al., 2015; Boyle et al., 2019).

Another striking feature of our optogenetic activation experiments was the widespread connectivity between iPVINs and unidentified neurons throughout laminae I-III. Many of these cells received direct inhibitory synaptic input from iPVINs, consistent with previous anatomical work showing axodendritic and axosomatic inputs from PVINs onto both PKCγ-expressing neurons and vertical cells in these laminae (Petitjean et al., 2015; Boyle et al., 2019). One other study has also used optogenetic activation of PVINs to assess inhibitory postsynaptic responses in spinal interneurons, and found that oIPSCs were observed in most recordings (Yang et al., 2015). Together, these observations have important functional implications, and demonstrate that postsynaptic inhibition mediated by iPVINs is likely to extend further than modulating activity of PKCγ-expressing and vertical cells. This widespread connectivity is also relevant to previous interpretations of work manipulating PVIN function in optogenetic and chemogenetic studies, as well as the impact of reduced iPVIN-mediated inhibition proposed as a substrate for tactile allodynia (Petitjean et al., 2015; Boyle et al. 2019).

### Neurotransmitter specialization at iPVIN inputs

Our data on iPVINs show these cells use both GABAergic and glycinergic signalling for postsynaptic inhibition, with a clear dominance of glycinergic input. These observations match our previous findings that PVINs themselves are inhibited by glycinergic mechanisms (Gradwell et al., 2017). This is also consistent with the view that GABAergic inhibition dominates in superficial lamina of the dorsal horn, whereas glycinergic inhibition is more important in deeper lamina (Graham et al., 2003; Cronin et al., 2004; Anderson et al., 2009; Graham et al., 2011; Zeilhofer et al., 2012). This strong glycinergic input from iPVINs in laminae IIi-III also has functional implications, given that the temporal properties of glycinergic and GABAergic currents differ in the superficial dorsal horn. Synaptic currents mediated by glycine have fast kinetics with a rapid decay time course (Lynch 2004), and this feature is thought to be important for the delivery of strong and precisely-timed inhibition in circuits involved in locomotor pattern generation (Callister and Graham 2010; Fink et al., 2014). The need for precision in inhibitory control of discrete circuits is equally important in the dorsal horn, and also favours the predominance of glycinergic inhibition to help segregate the passage of sensory information from functionally distinct classes of afferent fibres.

Contrasting the glycine dominance at postsynaptic iPVIN inputs, our data showed that iPVINs only utilize GABA at the axoaxonic inputs that mediate presynaptic inhibition. This is in line with a long literature demonstrating a critical role for GABA in mediating presynaptic inhibition (Eccles et al., 1963; Schmidt 1963; Davidoff 1972; Feltz and Rasminsky 1974; Gallagher et al., 1978) and excluding a role for glycine (Eccles et al., 1963; De Groat et al., 1972; Levy and Anderson 1972). This novel difference in the neurotransmitters for pre- and postsynaptic inhibition could be explained either by the existence of two specialised iPVIN populations, or specialisation at iPVIN terminals to determine the functional significance of GABA and glycine at different connections. The established pattern of co-expression of GABA and glycine in PVINs (Laing et al., 1994), along with the observations that most terminals presynaptic to myelinated primary afferents in the dorsal horn contain both GABA and glycine (Todd 1996), and the absence of glycine receptors in primary afferent terminals (Mitchell et al., 1993), supports the later explanation. Together, these results add to two decades of work revising Dale’s single transmitter principle for synaptic transmission (Pelky et al., 2020). Interestingly, these revisions began in the spinal cord with the demonstration of GABA/glycine co-transmission in the ventral spinal cord (Jonas et al., 1998) and iPVINs can be added as a source of such signals in the DH. Furthermore, our data emphasise iPVINs differentially use glycine and GABA at different locations across the circuits they regulate, with this functional selectivity optimised to efficiently segregate innocuous tactile input from pain-processing circuits.

### Role of ePVINs and iPVINs in pain processing

In addition to preventing low threshold input from activating nociceptive circuits (Petitjean et al., 2015; Boyle et al., 2019), our data show that both ePVIN and iPVIN populations are capable of modulating nociceptive circuits more directly. Previous anatomical assessment has shown PV axon terminals form a dense plexus within lamina IIi and spread into lamina III, whereas PV axon terminals are sparse in lamina IIo, and virtually absent in lamina I (Hughes et al. 2012, Petitjean et al. 2015). Despite this, our PVIN photostimulation experiments demonstrate ePVINs provide excitatory input to neurons in the superficial dorsal horn (laminae I-IIo). This discrepancy between anatomy and function suggests the majority of lamina I-IIo neurons receiving PVIN input, including projection neurons, do so via ventrally directed dendrites. Our observation that ePVIN input to these superficial populations was small amplitude and rarely evoked AP discharge in spinal cord slices is consistent with such inputs terminating on distal dendrites and undergoing electrotonic filtering. Our *in vivo* phototstimulation experiments show that when fully intact and strongly activated, ePVINs can recruit signalling in lamina I, observed as Fos activation in this region. In addition, several excitatory interneuron populations in lamina II are known to synapse with lamina I projection neurons including vertical cells (Lu and Perl., 2005; Cordero-Erausquin et al., 2009), calretinin-expressing interneurons (Smith et al., 2019; Petitjean et al., 2019) and CCK-expressing interneurons (Liu et al., 2018). Given that we now show that most ePVINs co-express CCK, and that ePVINs provide direct input to identified lamina I projection neurons, as well as presumptive excitatory interneurons in this region, our findings position these ePNINs as an element of this nociceptive circuitry. Finally, the high degree of inhibitory connections revealed by photostimulation suggests that this ePVIN-based nociceptive circuitry is under inhibitory regulation by iPVINs. Together, this builds on the established role of iPVINs as a ‘gate’ that prevents innocuous stimuli from accessing superficial nociceptive circuitry.

In conclusion, this work extends our understanding of the role PVINs play in spinal sensory circuits by mapping and manipulating the activity of both excitatory and inhibitory subtypes. We establish that there is a high degree of inhibitory connectivity between PVINs, however, we only detected anatomical evidence of excitatory interconnectivity. We show that ePVINs make connections throughout LI-III and activate nociceptive circuitry within LI-IIo, including projection neurons. These findings implicate ePVINs in the pathological recruitment of pain circuits following loss of PVIN inhibition, and add to those connections already established with lamina II vertical and PKCγ cells. Furthermore, iPVINs not only strongly inhibit deep tactile circuits through pre- and post-synaptic inhibition, but also contribute to the inhibition of superficial nociceptive circuitry within LI-IIo. Together, these findings are important for our understanding of PVINs, and how a diverse range of sensory modalities are appropriately encoded by neuronal circuits within the dorsal horn.

## Methods

### Animals

All procedures were approved by the Animal Care and Ethics Committee at the University of Newcastle. All experimental procedures performed at the University of Glasgow were conducted in accordance with the European Community directive 86/609/EEC and UK Animals (Scientific Procedures) Act 1986. All experiments were carried out on wild-type animals (C57Bl/6), PV^Cre^ knock-in mouse line (JAX Stock # 08069); or offspring of PV^Cre^ mice crossed with the Cre-dependent ChR2-YFP mouse line Ai32 (JAX Stock # 012569) or the Cre-dependent tdTomato reporter line Ai9 (JAX Stock # 012569).

### Immunocytochemistry in PV^Cre^;Ai9 mice

Three PV^Cre^;Ai9 mice (both sexes, 18-21 g) were perfused transcardially with fixative containing 4% formaldehyde in 0.1M phosphate buffer. Transverse sections (60 µm thick) of the lumbar enlargement (L3-L5) were cut on a vibrating microtome and immersed for 30 minutes in 50% alcohol to enhance antibody penetration. Sections were incubated for 3 days at 4°C with primary antibodies (See Key Resources Table), washed in double-salt phosphate-buffered saline (PBS), then incubated for a further 24 hours at 4°C with species-specific secondary antibodies raised in donkey. Sections were then washed again with double-salt PBS and mounted on glass coverslips in Vectashield anti-fade mounting medium (Vector Laboratories, Burlingame, CA). Primary and secondary antibodies were diluted in PBS containing 0.3% Triton X-100 and 5% normal donkey serum.

Tiled confocal stacks encompassing the entire section thickness of one dorsal horn were taken on three sections per animal on a Zeiss LSM 710 microscope system, using a 40x oil-immersion lens (NA = 1.3) and a z-step of 1 µm. Each channel of the resulting images (containing PV, Pax2 or mCherry immunoreactivity) was viewed separately and all cell bodies in laminae II_i_-III that were immuno-positive within each channel were marked throughout the section thickness using Neurolucida software (MBF Bioscience, Williston, VT). Cells were only included if the maximal profile of their soma lay within the section thickness. Once each channel had been assessed independently the marked cells were combined and co-expression of PV, Pax2 and/or mCherry was determined.

### Fluorescent in situ hybridization in wild-type mice

Three wild-type C57BL/6 mice (both sexes, 18–20□g) were used for *in situ* hybridization experiments, as described previously (Gutierrez-Mecinas et al., 2019). Animals were decapitated under deep isoflurane anaesthesia before their spinal cords were removed then rapidly frozen on dry ice. Fresh frozen lumbar spinal cord segments were embedded in OCT mounting medium then cut into 12 μm-thick transverse sections on a cryostat (Leica CM1860; Leica; Milton Keynes, UK) and mounted on SuperFrost Plus slides (48311[703; VWR; Lutterworth, UK). Multiple[labelling fluorescent *in situ* hybridization was performed with RNAscope probes and fluorescent multiplex reagent kit 320850 (ACD BioTechne; Newark, CA). Reactions were carried out according to the manufacturer’s recommended protocol. Probes used in this study were GAD1 (Cat no. 400951), Slc17a6 (Cat no. 319171), PValb (Cat no. 421931) and CCK (Cat no. 402271). Sections were reacted with the probe combinations GAD1, Slc17a6 and PValb or CCK, Slc17a6 and PValb, revealed with Atto 550, Alexa 647, and Alexa 488, respectively. Sections were mounted in Prolong-Glass anti-fade medium with NucBlue (Hoechst 33342; ThermoFisher Scientific, Paisley, UK). RNAscope positive and negative control probes were tested on other sections simultaneously.

Three sections per animal were selected prior to viewing *in situ* hybridization florescence (to avoid bias) and imaged using the 40x oil-immersion lens with the confocal aperture set to 1 Airy unit. In each case, tile scanning of a single optical plane through the middle of the section was used to include the whole of laminae I-III. Semi-automated analysis of transcript numbers per nucleus was conducted using the cell detection and subcellular objects features on Qupath software (Bankhead et al., 2017). Cell analysis was conducted only in laminae II_i_ and III, where a band of dense parvalbumin transcripts were present. Recognition and segmentation of individual nuclei was performed based on NucBlue staining. An additional 2 μm perimeter was added to each nucleus to allow detection of perinuclear transcripts. This additional perimeter was omitted where cells were directly adjacent to each other. Any areas with poor nuclear segmentation were excluded manually from the analysis following examination of each segmented section. Single RNA transcripts for each target gene appeared as individual puncta and detection thresholds were adjusted manually until the mark-up accurately reflected the transcript distribution. Data output consisted of manual inspection of the section to ensure accuracy, followed by export of a table containing each cell’s transcript numbers. This was further analysed in Microsoft Excel. Cells were defined as positive for expression of a given gene if they contained greater than four transcripts. Cells were classified as excitatory or inhibitory depending on expression of Slc17a6 or GAD1, respectively.

### Intraspinal Injection of Brainbow-encoding AAVs

To visualise the somatodendritic arbors of individual PV-expressing spinal interneurons, we performed co-injections of two adeno-associated viruses (AAVs) carrying Cre-dependent Brainbow expression cassettes in PV^Cre^ mice (n = 5; both sexes; 18-23 g at surgery): AAV.Brainbow1 codes for eYFP and TagBFP, whereas AAV.Brainbow2 codes for mTFP and mCherry (Cai et al., 2013). By adopting this approach, individual cells displayed a unique colour profile based on the stochastic expression of farnesylated fluorescent proteins encoded by the AAVs.

To perform these intraspinal AAV injections, animals were anaesthetised with isoflurane (5% induction, 1.5 - 2% maintenance) and placed in a stereotaxic frame. Bilateral injections were made into the L3 dorsal horns using a glass micropipette attached to a 10 µL Hamilton syringe. Injections were made through the T12/13 intervertebral space, 400 µm lateral to the midline and 300 µm below the pial surface. For each injection, 500 nL of virus was infused at a rate of 30-40 nL/minute using a syringe pump (Harvard Apparatus, MA, USA). Given that the aim of this experiment was to reconstruct the morphology of individual PV-expressing cells, we used moderate titres of viruses (3.77×10^8^ GC for AAV.Brainbow1 and 3.72×10^8^ GC for AAV.Brainbow2) to achieve sparse labelling of PV neurons. By labelling only a small proportion of PV neurons at random, this ensures that the morphology of individual neurons can be traced with confidence. Once injections were complete, wounds were closed and animals allowed to recover with appropriate analgesic administration (0.3mg/kg buprenorphine and 5mg/kg carprofen). All animals made uneventful recoveries from the intraspinal injection surgery.

### Morphological analyses of Brainbow-labelled PV interneurons

Sagittal sections (60 µm thick) of the L3 spinal segment were processed for immunocytochemistry as detailed above. For the analysis of neuronal morphology, sections were incubated in a cocktail of primary antibodies to reveal three (tagBFP, mTFP and mCherry) of the four fluorescent proteins as well as Pax2 expression to label inhibitory interneurons (Foster et al., 2015; Larsson, 2017). Tiled confocal scans of Brainbow-labelled cells within laminae IIi-III were made through the full thickness of the sections using the 40x objective (1.5x zoom and 0.5µm z-step). Cells were selected for reconstruction if: 1) they demonstrated relatively strong staining for at least one fluorescent protein (to allow them to be readily distinguished from neighbouring labelled cells based on colour), and 2) their soma was found near the middle of the section in the z-axis (to ensure maximal representation of their dendritic arbor in the medio-lateral plane). The presence or absence of Pax2 immunolabelling in these cells was then determined. Based on these selection criteria, the somato-dendritic morphology of 30 inhibitory (Pax2-expressing) and 34 excitatory (Pax2-lacking) Brainbow-labelled PV interneurons were reconstructed in three dimensions using Neurolucida software (see Table 2). To compare the morphology of inhibitory and excitatory PV interneurons objectively, fifty morphological parameters (5 for the soma and 45 for the dendritic arbor) were extracted from the Neurolucida reconstructions (Dickie et al., 2019; Iwagaki et al., 2016) using Neurolucida Explorer (MBF Bioscience, Williston, VT). K-means non-hierarchical clustering was performed in Orange software (University of Ljubljana) based on these 50 morphological parameters, with the number of clusters set to 2 and cluster seeds chosen using the K-means++ algorithm. The 3D graph displaying the relationship of 3 of the morphological variables within the K-means-derived clusters shown in Figure 2F was produced in TeraPlot (Kylebank Software Ltd., Ayr, UK).

**Table 2.**
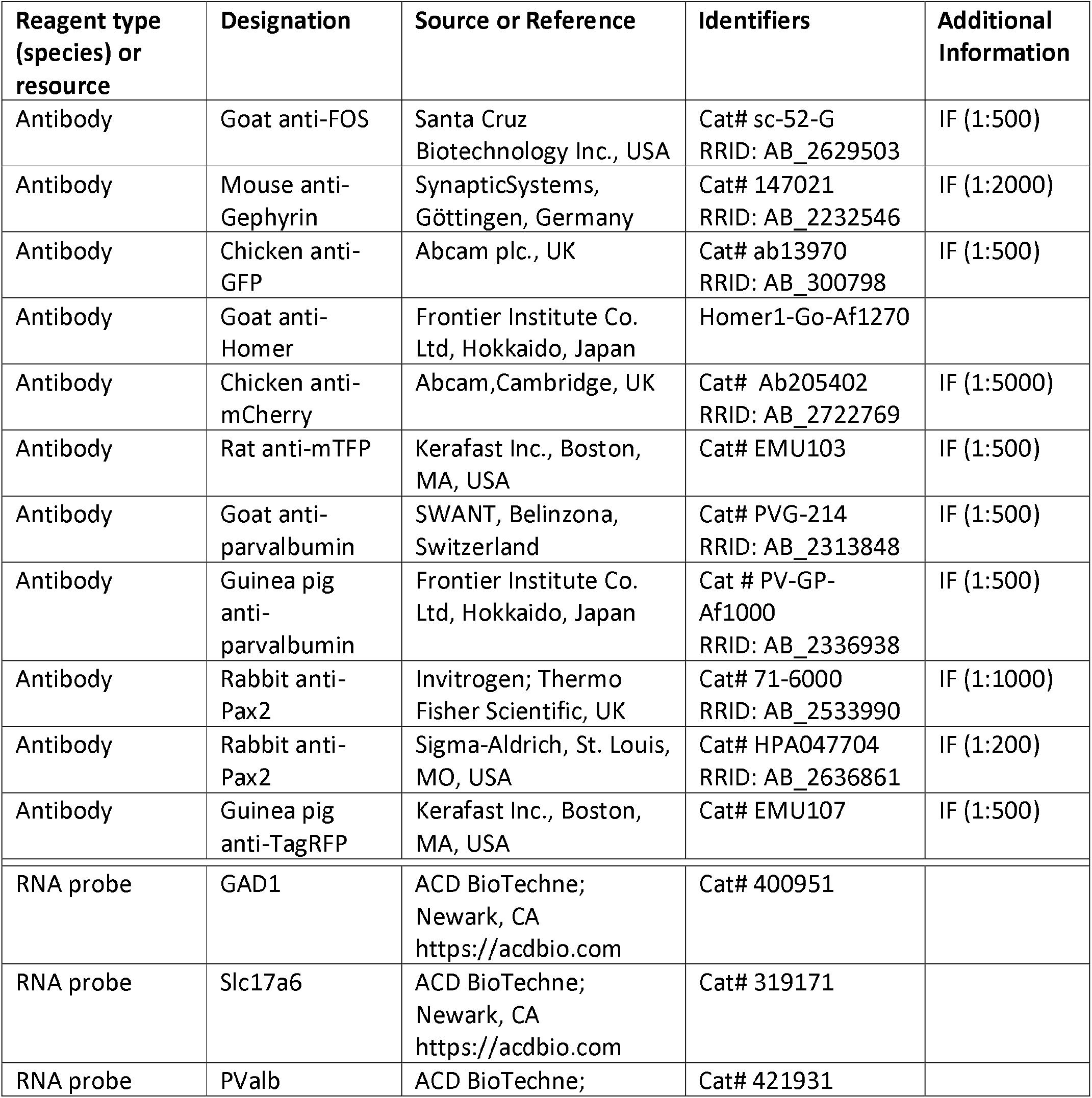
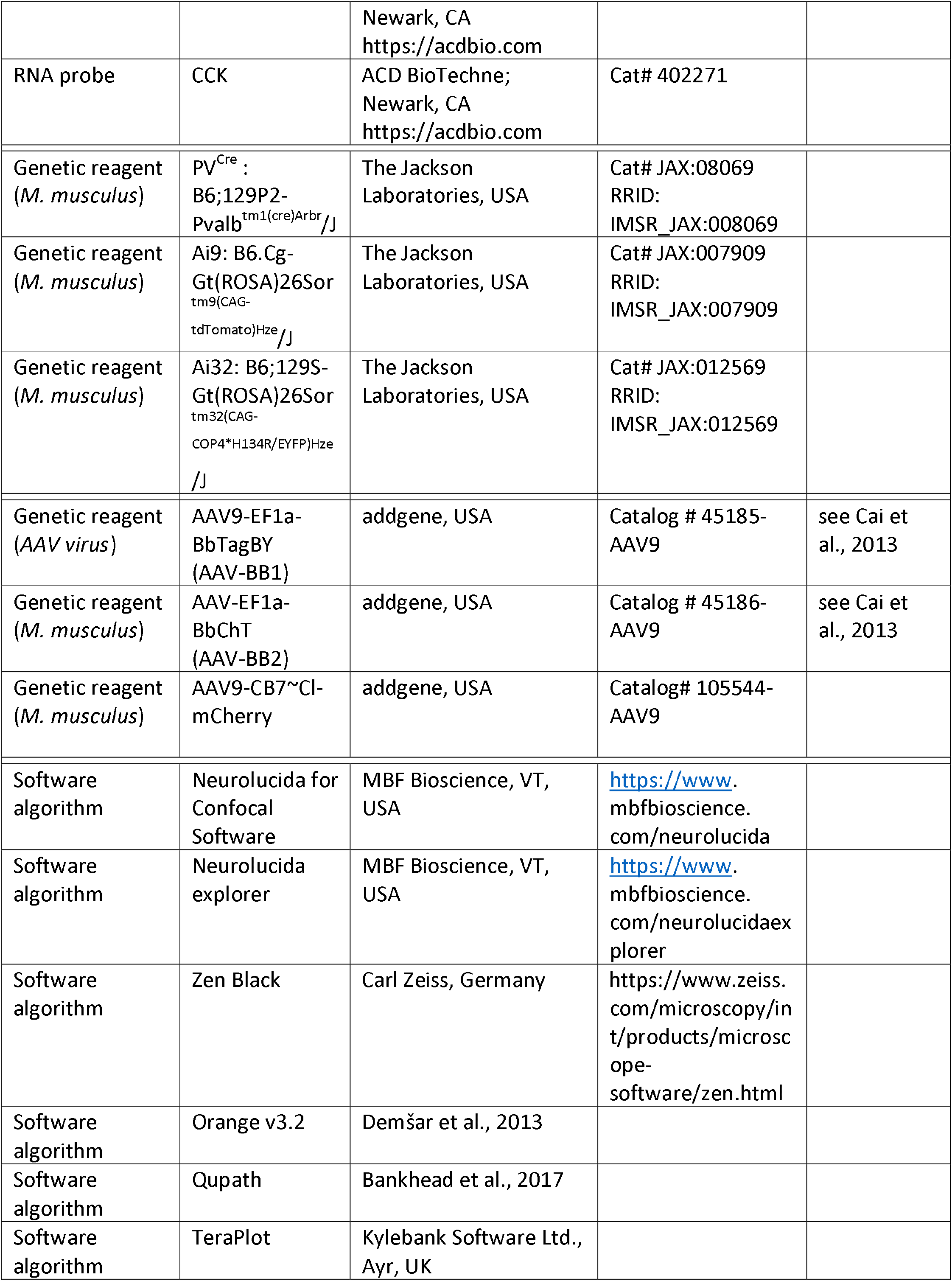
Resources and reagents.

To determine whether excitatory and inhibitory PV interneurons form homotypic and/or heterotypic synaptic circuits between themselves, we used antibodies to reveal two of the Brainbow fluorescent proteins (mTFP and mCherry) combined with immunolabelling for gephyrin and Homer1 to label inhibitory and excitatory synapses, respectively. Given that we were limited in the number of flurophores we were able to stain for in our experiment (total of 4), we stained Pax2 immunoreactivity in the same channel as that for Homer1, since the two proteins can be easily differentiated based on their sub-cellular localisation (i.e. Pax2 is expressed in the nucleus, whereas Homer1 produces punctate membrane labelling).

### Electrophysiology

All electrophysiological studies were performed on PV^Cre^;Ai32 mice (both sexes; age = 30 ± 3 weeks). Mice were deeply anesthetized with ketamine (100 mg/kg, i.p.) and decapitated. Using a ventral approach, the lumbar enlargement of the spinal cord was rapidly removed and glued to the stage of a vibrating microtome (Leica VT-1000S, Heidelberg, Germany). Sagittal or transverse slices (200 µm thick) were prepared in a bath of carbogenated ice-cold sucrose-substituted ACSF containing (in mM): 250 sucrose, 25 NaHCO_3_, 10 glucose, 2.5 KCl, 1 NaH_2_PO_4_, 1 MgCl_2_ and 2.5 CaCl_2_. All slices were incubated for 1 hr at 22-24°C in an interface chamber containing carbogenated ACSF (118 mM NaCl substituted for sucrose) before recording commenced.

Slices were transferred to a recording chamber and continually superfused (bath volume 0.4 ml; exchange rate 4–6 bath volumes/min.) with ACSF bubbled with Carbogen (95% O_2_ and 5% CO_2_) to achieve a pH of 7.3–7.4. Recordings were obtained at room temperature (21–24°C) and neurons were visualized with a Nikon FN-PT with near-infrared differential interference contrast optics connected to a camera (Jenoptik ProgRes MF cool). Recordings were made in two locations: 1) within the clearly discernible YFP-expressing plexus of PVINs (laminae II_i_-III); and 2) superficial to the PVIN plexus (laminae I-II_o_). Furthermore, three neuron types were targeted in recordings 1) PVINs identified by YFP expression and ChR2-mediated photocurrents; 2) unidentified dorsal horn neurons that lacked YFP and did not exhibit photocurrents; and 3) virally-labelled projection neurons (PNs). Slices were illuminated using a CoolLED pE excitation system that allowed: visualization of YFP fluorescence, ChR2 photostimulation using a FITC filter set, and visualization of mCherry-expressing PNs with TRITC filters.

Recordings were acquired in voltage-clamp (holding potential −70 mV) or current clamp (maintained at −60 mV using small (< 20 pA) bias current injection. Patch pipettes (4-8 MΩ) were filled with a cesium chloride-based internal solution containing (in mM): 130 CsCl, 10 Hepes, 10 EGTA, 1 MgCl_2_, 2 ATP and 0.3 GTP (pH adjusted to 7.35 with 1 M CsOH) for assessing inhibitory synaptic transmission. A potassium gluconate-based internal solution containing (in mM): 135 C6H11KO7, 6 NaCl, 2 MgCl2, 10 HEPES, 0.1 EGTA, 2 MgATP, 0.3 NaGTP, pH 7.3 (with KOH) was used to record AP discharge or excitatory synaptic transmission (internal solution osmolarity adjusted to 280 and 300 mOsm, respectively). In recordings from YFP-expressing neurons, QX-314 bromide was added to the internal solution to block fast activating voltage-gated sodium channels. This avoided contamination of recordings with unclamped spikes. Neurobiotin (0.2%) was included in both internal solutions for post-hoc confirmation of neuronal morphology (Vector Laboratories, Burlingame, CA, USA).

Data were amplified using a Multiclamp 700B amplifier (Molecular Devices, Sunnyvale, CA, USA) digitized online (sampled at 10–20 kHz and filtered at 5–10 kHz) via an ITC-18 computer interface (Instrutech, Long Island, NY, USA), acquired and stored using Axograph X software (Axograph X, Sydney). After obtaining the whole-cell recording configuration, series resistance, input resistance and membrane capacitance were calculated based on the response to a 5 mV hyperpolarising voltage step (10 ms duration) from a holding potential of −70 mV. These values were monitored at the beginning and end of each recording session and data were rejected if values changed by more than 30%.

Photostimulation intensity was suprathreshold (16 mW) with a duration of 1 ms (controlled by transistor-transistor logic pulses), unless otherwise stated. This ensured generation of action potential discharge in PV-ChR2/YFP neurons and allowed confident assessment of postsynaptic currents in recorded neurons. To isolate monosynaptic connectivity, 1μM TTX and 200μM 4-AP were bath-applied to block action potential discharge and accentuate light-evoked neurotransmitter release from ChR2-expressing terminals (Petreanu et al. 2009, Cruikshank et al. 2010). To assess the impact of PV^+^IN activation on action potential discharge responses in unidentified neurons (i.e. cells lacking YFP) a series of three depolarizing step-currents were repeated (1000 ms or 50 ms duration, 20 pA increments, delivered every 10 seconds). During this protocol PV^+^INs were activated by photostimulation (16 mW, 10 ms) at the onset of second series of depolarizing steps and AP discharge was compared with the preceding and subsequent responses.

To identify lamina I projection neurons in slice recording experiments, a subset of animals (n=8) underwent surgery to inject a viral tracer, specifically AAV9.CB7.CI.mCherry (viral titre = 2.5 x1013 vg/mL), into the parabrachial nucleus (PBN) to retrogradely-label lamina I PNs for subsequent targeted patch-clamp recording experiments. Briefly, mice were anaesthetised with isoflurane (5% induction, 1.5-2% maintenance) and secured in a stereotaxic frame (Harvard Apparatus, Massachusetts, U.S.A). Two small craniotomies were performed and up to 700 nL of viral sample was injected using a picospritzer (PV820, WPI, Florida, USA) into the PBN bilaterally. These injections were made 5.25 mm posterior to bregma, ± 1.2 mm of midline and 3.8 mm deep from skull surface, using coordinates refined from those in the mouse brain atlas (Paxinos and Franklin, 2001).

Injections were made over 5 minutes and the pipette was left in place for a further 7-10 minutes to avoid drawing the virus sample along the pipette track. Animals recovered for 2-4 weeks to allow sufficient retrograde labelling of projection neurons before spinal cord slices were prepared. Spinal cord slices were obtained using methods described above (Electrophysiology) and mCherry positive neurons were visualised for recording using a Texas Red filter set. PVIN photostimulation was performed as above for other DH populations. The brain from each animal was also removed and brainstem sectioned slices, containing the PBN, were prepared to confirm the injection site. In all cases the injection site was focussed on the PBN.

### Optogenetic stimulation for Fos activation mapping

The postsynaptic circuits targeted by PVINs were assessed by delivering spinal photostimulation to anaesthetised PV^Cre^;Ai32 animals and then processing spinal cords for Fos-protein as described previously (Smith et al., 2019). Animals (n = 3) were anaesthetised with isoflurane (5% initial, 1.5-2% maintenance) and secured in a stereotaxic frame. A longitudinal incision was made over the T10-L1 vertebrae and a laminectomy was performed on the T13 vertebra. Unilateral photostimulation (10 mW, 10 ms pulses @ 10 Hz for 10 min) was then delivered to the exposed spinal cord by positioning an optic fiber probe (400 nm core, 1 mm fiber length, Thor Labs, New Jersey, U.S.A) above the spinal cord surface using the stereotaxic frame. Photostimulation was delivered by a high intensity LED light source attached to the probe via a patch cord. Following photostimulation animals remained under anaesthesia for a further 2 hrs for subsequent comparison of Fos expression in the dorsal horn. Animals were then anaesthetised with ketamine (100 mg/kg i.p.) and perfused transcardially with saline followed by 4% depolymerised formaldehyde in 0.1M phosphate buffer. Sections were processed for immunocytochemistry by incubating in a cocktail of antibodies including chicken anti-GFP and goat anti-Fos. Primary antibody labelling was detected using species-specific secondary antibodies conjugated to rhodamine and Alexa 488 (Jackson Immunoresearch, West Grove, PA, USA).

### Electrophysiology data analysis

All data were analysed offline using Axograph X software (Axograph X, Sydney). AP discharge was classified according to previously published criteria (Graham et al. 2004, Graham et al. 2007). Criterion for inclusion of a neuron for analysis was an RMP more negative than –50 mV and a series resistance < 30 MΩ (filtered at 5 KHz). In the analysis of AP discharge, individual APs elicited by step-current injection were captured using a derivative threshold method (dV/dt > 15 V/s) with the inflection point during spike initiation defined as AP threshold. AP amplitude was defined as the difference between AP threshold and the maximum positive peak associated with a spike. AP base-width was measured at AP threshold. AP afterhyperpolarization (AHP) amplitude was taken as the difference between AP threshold and the maximum negative peak following the spike. Rheobase current was defined as the smallest current step that elicited at least one AP, and AP latency was measured as the time difference between the stimulus onset (current injection or photostimulation) and AP threshold.

Most data assessed optically evoked excitatory postsynaptic currents (oEPSCs) and inhibitory postsynaptic currents (oIPSCs) in recorded neurons during PVIN photostimulation. When this analysis was undertaken in targeted PVIN recordings, an immediate photocurrent was also directly evoked during photostimulation followed by synaptic responses. In these instances, the baseline current was set to zero at the onset of the synaptic response, with the amplitude of photostimulation-evoked synaptic input then measured from this level. In all other recordings (i.e. cells lacking YFP) synaptic responses were measured from baseline just prior to photostimulation. The peak amplitude of response was calculated from the average of 10 successive trials. A number of parameters were considered for determining whether a photostimulation-evoked synaptic input was mono- or polysynaptic. The latency of oPSCs was measured as the time from photostimulation to the beginning of the evoked current. The ‘jitter’ in latency was measured as the standard deviation in latency of 10 successive trials. Importantly the latency of monosynaptic inputs was much shorter, there was minimal jitter in the onset of responses between trials, and reliability (percentage of photostimulation trials to evoke a response) was higher than those deemed to be polysynaptic inputs. To assess the contribution of different neurotransmitter systems to photostimulation responses neurotransmission blockers were applied. Changes in photostimulation-evoked postsynaptic current amplitude were measured to calculate either an oIPSC_index_ or oEPSC_index_ (Fink et al., 2014). These were calculated using the amplitude of the oPSC (oPSCλΦ) in the presence of the drug, and amplitude of oPSCs (oPSCλ) at baseline (i.e. no drug) in the following formula:

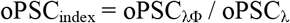

The value of the oPSC_index_ is 1 when the drug has no effect and 0 when the drug completely blocks the oPSC. Note, an oPSC_index_ of 0 was not possible for cells lacking YFP due to small variations in baseline noise, however, photocurrent zeroing prior to photostimulation-evoked oPSCs in PVIN recordings resulted in an oPSC_index_ of 0 under drug block conditions.

### Statistics

Statistical analysis was carried out using SPSS v10 (SPSS Inc. Chicago, IL, USA) and Prism 7 (GraphPad Software Inc., San Diego, CA, USA). Student t-tests and student-Newman-Keuls ANOVAs were used to compare variables. Data that failed tests for normal distribution and homogeneity of variance were compared using the non-parametric Kruskal-Wallace or Mann-Whitney tests. Statistical significance was set at p < 0.05. All values are presented as means ± SEM unless otherwise stated.

## LEAD CONTACT AND MATERIALS AVAILABILITY

Further information and requests for resources and reagents should be directed to and will be fulfilled by the Lead Contacts, Dr. Brett A. Graham or David I. Hughes.

---

**Figure S1 (Relates to Figures 2 and 3). Fidelity of tdTOM labelling in different CNS regions in the PV^**Cre**^;Ai9 mouse line.** The colocalisation of tdTom-expression and immunolabelling for PV was assessed in several different regions of the CNS. **A**, The general distribution of tdTOM cells in the lumbar spinal cord matched the general distribution of PV-IR cells, with a dense plexus of cells in laminae IIi and III, numerous cells in lamina V, and more scattered cells in ventral horn laminae VII and VIII. **B**, Most tdTOM cells in laminae IIi and III (asterisk) co-expressed PV-IR, although several PV-IR cells showed no labelling for tdTOM (arrows). **C-F**, The fidelity of tdTOM expression in PV-IR cells was much higher in the ventral horn (**C**), CA1 of the hippocampus (**D**), the dentate gyrus (**E**) and the cerebellum (**F**) with high incidence of co-expression of tdTOM labelling (red) in PV-IR cells (green), and PV-IR in tdTOM cells. Abbreviations panel D: S.O. = *stratum oriens*, S.P. = *stratum pyramidale*, S.R. = *stratum radiatum*. Abbreviations panel E: Mol = molecular layer, GCL =granule cell layer, Pol = polymorphic layer. Abbreviations panel F: Mol = molecular layer, PCL = Purkinje cell layer, GL = granular cell layer. Scale bars (µm): A = 250; B = 20; C = 50; D-F = 100.

