## Supplementary figures and images for "Three neurotransmitters regulate diverse inhibitory and excitatory Parvalbumin interneuron circuits in the dorsal horn"

### Supplemental Figure S1

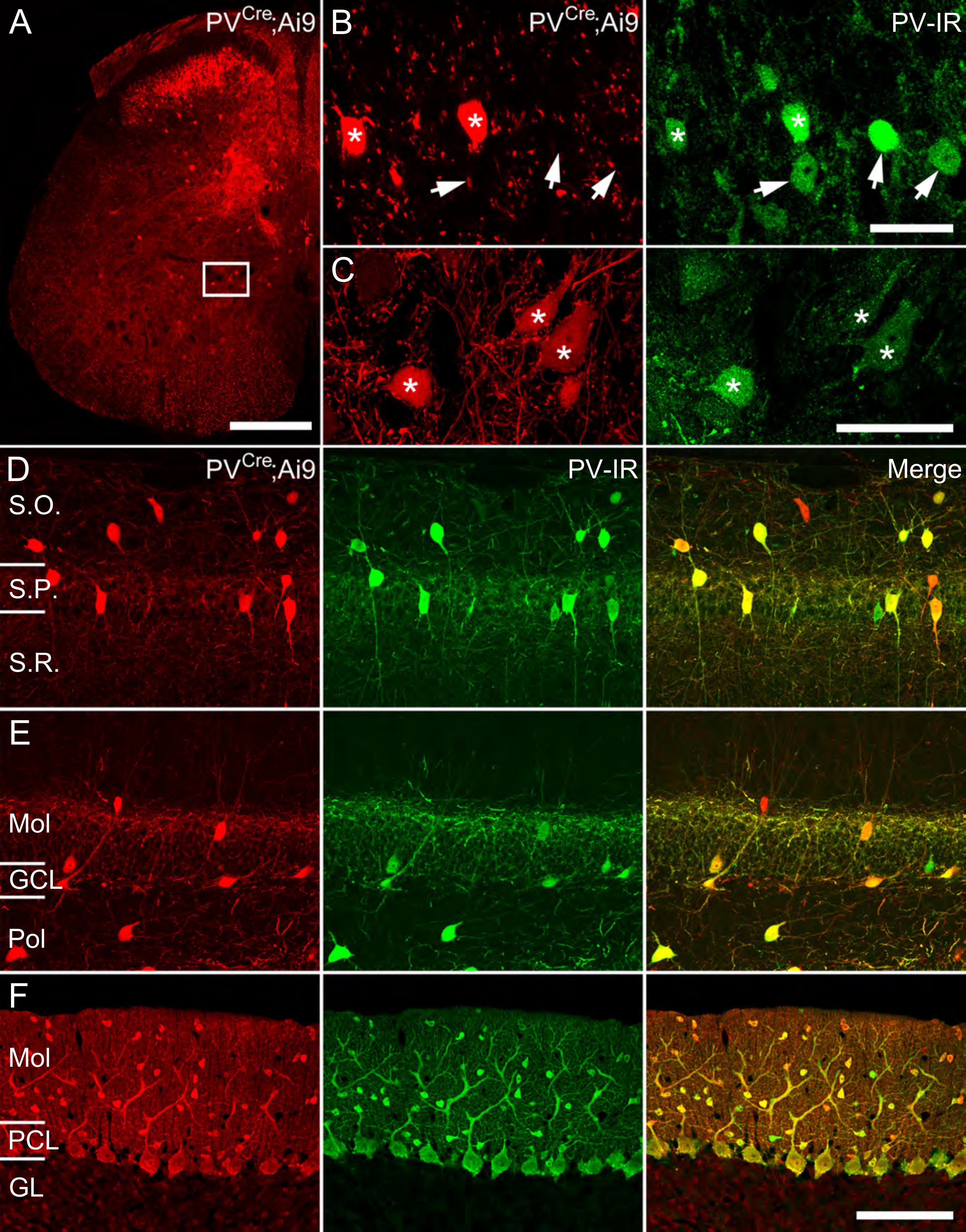
